# CCR7^+^ DC Define a Type 17 Module in Psoriasis

**DOI:** 10.1101/2024.04.28.591489

**Authors:** Yang Sun, Fangzhou Lou, Xiaojie Cai, Zhikai Wang, Xiuli Yang, Libo Sun, Zhouwei Wu, Zhaoyuan Liu, Yu-Ling Shi, Florent Ginhoux, Honglin Wang

**Author notes:** Correspondence to: Honglin WANG, Ph.D. Precision Research Center for Refractory Diseases, Shanghai General Hospital, Shanghai Jiao Tong University School of Medicine, Shanghai 201620, China. These authors contributed equally (Yang Sun, Fangzhou Lou).

## Abstract

Interleukin (IL)-23 is the master pathogenic cytokine in psoriasis and neutralization of IL-23 alleviates psoriasis. Psoriasis relapses after the withdrawal of IL-23 antibodies, and the persistence of IL-23-producing cells probably contributes to such recurrence. However, the cellular source of IL-23 was unclear, which hinders the development of targeted therapies focusing on modulating IL-23 expression aimed at resolving relapse. Here, we showed that *IL4I1*^+^*CD200*^+^*CCR7*^+^ dendritic cells (CCR7^+^ DC) dominantly produced IL-23 by concomitantly expressing the IL-23A and IL-12B subunits in human psoriatic skin. Deletion of CCR7^+^ DC completely abrogated IL-23 production in a mouse model of psoriasis and enforced expression of IL-23a in CCR7^+^ DC elicited not only αβT cell-driven psoriasis-like skin disease, but also arthritis. CCR7^+^ DC co-localized with CD161^+^ IL-17-producing T cells and KRT17^+^ keratinocytes, which were located in the outermost layers of psoriatic epidermis and strongly exhibited IL-17 downstream signatures. Based on these data, we identified CCR7^+^ DC as the source of IL-23 in psoriasis, which paves the way for the design of therapies focused on manipulating IL-23 production that may resolve the relapse of chronic inflammatory disorders like psoriasis.

**HIGHLIGHTS:**

- *IL4I1*^+^*CD200*^+^*CCR7*^+^ DC are dominant IL-23 producers in psoriasis and its mouse model.
- Psoriatic CCR7^+^ DC likely arise from cDC2.
- CD161 marks all IL-17-producing T cells in psoriatic skin.
- IL-23a overexpression in CCR7^+^ DC elicits an αβT cell-driven mouse model of psoriasis and arthritis.
- CCR7^+^ DC spatially define a type 17 module in psoriatic epidermis.

## INTRODUCTION

Psoriasis is a chronic and recurrent immune-mediated skin disorder affecting 125 million people globally(Armstrong and Read, 2020). It manifests as erythematous skin plaques covered with white scales, which impact the quality of life of patients both physically and psychologically(Griffiths and Barker, 2007). Although neutralization of pro-inflammatory cytokines, including IL-17A and IL-23, alleviates psoriasis symptoms, many patients are unresponsive or respond only partially to such therapies. In addition, responsive patients face the risk of drug resistance, or recurrence following the discontinuation of biologic agents(Amschler et al., 2020; Bito et al., 2014; Galluzzo et al., 2018; Masson Regnault et al., 2022), the latter being a key challenge in psoriasis therapy(Puig et al., 2022).

Interleukin (IL)-17-producing CD4^+^ T helper 17 (Th17) cells were initially identified as crucial players in psoriasis-associated-immune circuits. Later, CD8^+^ T cells were found to contain the T cytotoxic 17 (Tc17) population, giving rise to *in situ* IL-17 production(Kryczek et al., 2008; Lowes et al., 2008; Ortega et al., 2009). Recent studies revealed key roles of CD8^+^ tissue-resident memory T (T_RM_) cells in the epidermis as sources of IL-17 in inflamed psoriatic skin, and as rapid responders in recall responses in resolved psoriasis(Cheuk et al., 2017; Cheuk et al., 2014). These T_RM_ cells are considered to be the immediate cause of the relapse of psoriasis(Conrad et al., 2007). While IL-17 is produced mainly by αβT cells in human psoriatic skin, psoriasis-like mouse models are mostly driven by γδT cell-derived IL-17(Cai et al., 2011; Mabuchi et al., 2011). This reflects distinct differences in the immunopathological principles in human psoriasis versus mouse models, which hinders the effective evaluation of drug candidates aimed at preventing psoriasis recurrence.

As subpopulations of the dendritic cell (DC) population, both plasmacytoid DCs (pDC) and conventional DCs (cDC), are considered drivers of psoriasis. The type 1 IFN-producing pDC are crucial for the initiation of psoriasis(Nestle et al., 2005), while the cDC secrete IL-23 as a third signal that induces polarization toward a type 17 T cell response in the disease(Wohn et al., 2013). Specific targeting of IL-23 in psoriasis has been demonstrated to effectively prolong the time to relapse, which further emphasizes the key role of IL-23 in the recurrence and maintenance of psoriasis, and highlights manipulation of IL-23 production as a potential therapeutic strategy(Masson Regnault et al., 2022). Although IL-23 in psoriasis-like mouse skin is reported to be produced by CD301b^+^ cDC2, the counterpart of these DCs in human skin as well as the exact cellular source of IL-23 in psoriasis remain elusive(Guttman-Yassky et al., 2011; Hansel et al., 2011; Kim et al., 2018; Nakamizo et al., 2021; Whitley et al., 2022). Thus, identification of the IL-23 producers in psoriasis is a critical step for developing novel therapeutic strategies aimed at resolving relapse.

Here, we showed that *IL4I1*^+^*CD200*^+^*CCR7^+^* DC (CCR7^+^ DC), but not other myeloid cells, produced IL-23 by concomitantly expressing the IL-23A and IL-12B subunits, both of which are required to form the intact IL-23 for extracellular secretion, in psoriasis. Furthermore, we identified CCR7^+^ DC as the major source of IL-23 in the skin in the imiquimod (IMQ)-induced psoriasis-like mouse model. Importantly, Il4i1^+^ cell-specific overexpression of IL-23a elicited an αβT cell-driven psoriasis-like mouse model, which also developed psoriatic arthritis-like symptoms, with lesions that transcriptionally mimicked human psoriasis. In human psoriatic skin, IL-23-producing CCR7^+^ DC localized spatially with IL-17-producing CD161^+^ T cells and the IL-17-responsive KRT17^+^ keratinocyte subpopulation, defining a type 17 module that can be harnessed for the discovery of drug targets and rational design of therapeutic strategies for psoriasis.

## RESULTS

### Psoriatic keratinocytes exhibit a reconstructed differentiation trajectory

To identify immune and non-immune cells that might be involved in the development of psoriasis, we generated transcriptomes of individual epidermis and dermis cells obtained from six psoriasis patients and four healthy donors using the 10× Genomics platform (**Figures S1A** and **S1B**). After the epidermis was dissociated from the dermis by enzymatic digestion, live epidermal cells and live CD45^+^ dermal leukocytes were sorted from each sample using fluorescence activated cell sorting (FACS) and subjected to 3’-barcoded scRNA-seq to generate unique molecular identifier (UMI) counts matrixes (**Figures S1C** and **S1D**). After quality control and doublet exclusion, 31,750 epidermal cells and 42,054 dermal leukocytes from the 20 samples were integrated and clustered jointly. We performed uniform manifold approximation and projection (UMAP) dimensional reduction and partitioned the cells according to their respective marker genes (**Figures S1D** and **S1E**)(Becht et al., 2018). We identified keratinocytes, T cells, myeloid cells, mast cells, pDC and B cells in our data (**Figures S1D** and **S1E**).

To characterize non-immune cells in psoriatic epidermis, we extracted keratinocytes from the integrated data of all 10 epidermal samples for UMAP dimensional reduction and named the sub-clusters according to their hallmark genes (**Figures S2A** and **S2B**). Comparison of the interfollicular keratinocyte sub-clusters of psoriatic epidermis to those of healthy epidermis showed that: (1) *KRT14* and *KRT5* were highly upregulated not only in basal keratinocyte sub-clusters (KRT14^+^_ASS1^+^ and KRT14^+^_KRT15^hi^), but also in suprabasal keratinocyte sub-clusters of psoriatic epidermis; (2) spinous keratinocytes (KRT10^+^_KRT5^hi^ and KRT10^+^_KRT2^+^) normally express *KRT10*, *KRT1* and *KRT2*, while these genes were downregulated in psoriatic lesions; (3) Expression of *KRT6*, *KRT16* and *KRT17*, which were shown to be induced by T cell-derived cytokines including IFN-γ, IL-17 and IL-22(Yang et al., 2017; Zhang et al., 2019), was higher in psoriatic keratinocytes than in normal controls, with upregulated *KRT17* specifically detected in *KRT17*^+^ keratinocytes (KRT17^+^) and granular keratinocytes (FLG^+^) of psoriatic epidermis; and (4) psoriatic keratinocytes upregulated IL-17A-downstream genes including *S100A8*, *S100A9*, *SERPINB3* and *SERPINB4*, and IFN-γ-downstream genes including *ISG15*, *IFITM1*, *IFI6*, *IFITM3* and *IFI27*, in comparison to healthy keratinocytes (**Figures S2B** and **S2C**). Employing a pseudo-time trajectory to potentially understand the differentiation programs of healthy and psoriatic keratinocytes, we found a delayed decline in *KRT14* expression, abnormal upregulation of *MKI67* and *KRT17* in the end of the pseudo-time line, and the absence of the terminal differentiation marker *LOR* in psoriatic interfollicular keratinocytes (**Figure S2D**). These data highlighted dysregulated keratinocyte differentiation and proliferation in the different layers of psoriatic epidermis, a process that is probably regulated by diverse inflammatory cues from T cells.

### KRT17^+^ keratinocytes respond to IL-17 derived from CD161^+^ T cells

To gain functional insights into the T cells driving such aberrant keratinocyte differentiation, we investigated the T cell heterogeneity in psoriatic skin in comparison to normal skin. *CD3*^+^ T cells were extracted from the integrated data, and further sub-clustered (**Figures 1A** and **1B**). *KLRB1*^+^ (CD161^+^) T cells (KLRB1_T) and *CD8*^+^ T cells (CD8_Cytotoxic_T and CD8_T_RM_) expressed high levels of T_RM_ cell markers including *CD69* and *ITGAE*, while *CD4*^+^*FOXP3*^+^ T_reg_ cells expressed *CCR7*, but not T_RM_ cell markers (**Figure S3A**). Importantly, *IL17A*, *IL17F* and *IL26* expression was almost completely confined to “KLRB1_T” cells (**Figure 1B**), which were characterized as *KLRB1*^+^ αβT cells as these cells expressed *TRAC*, but not *TRDC* (**Figure S3A**). Notably, increased percentages of *IL17A*^+^ T cells, *IL17F*^+^ T cells and *IL26*^+^ T cells were identified among the *KLRB1*^+^ αβT cells of psoriatic skin compared to normal skin (**Figure 1C**). Given that CD4^+^ Th17 cells and CD8^+^ Tc17 cells were reported to produce IL-17 in psoriasis(Ho and Kupper, 2019), we calculated the ratios of *CD4*^+^, *CD8*^+^, *CD4*^+^*CD8*^+^ and *CD4*^-^*CD8*^-^ among the *KLRB1*^+^ αβT cells. The *KLRB1*^+^ αβT cell population consisted of approximately 20% *CD4*^+^ T cells, 20% *CD8*^+^ T cells and 60% *CD4*^-^*CD8*^-^ T cells, and these percentages did not differ between normal and psoriatic skin samples (**Figure S3B**). Notably, a significantly higher proportion of the *CD4*^-^*CD8*^-^ T cells expressing *IL17A* among the *KLRB1*^+^ αβT cell population was found in psoriatic skin compared to normal skin (**Figure 1D**).

**Figure 1.**
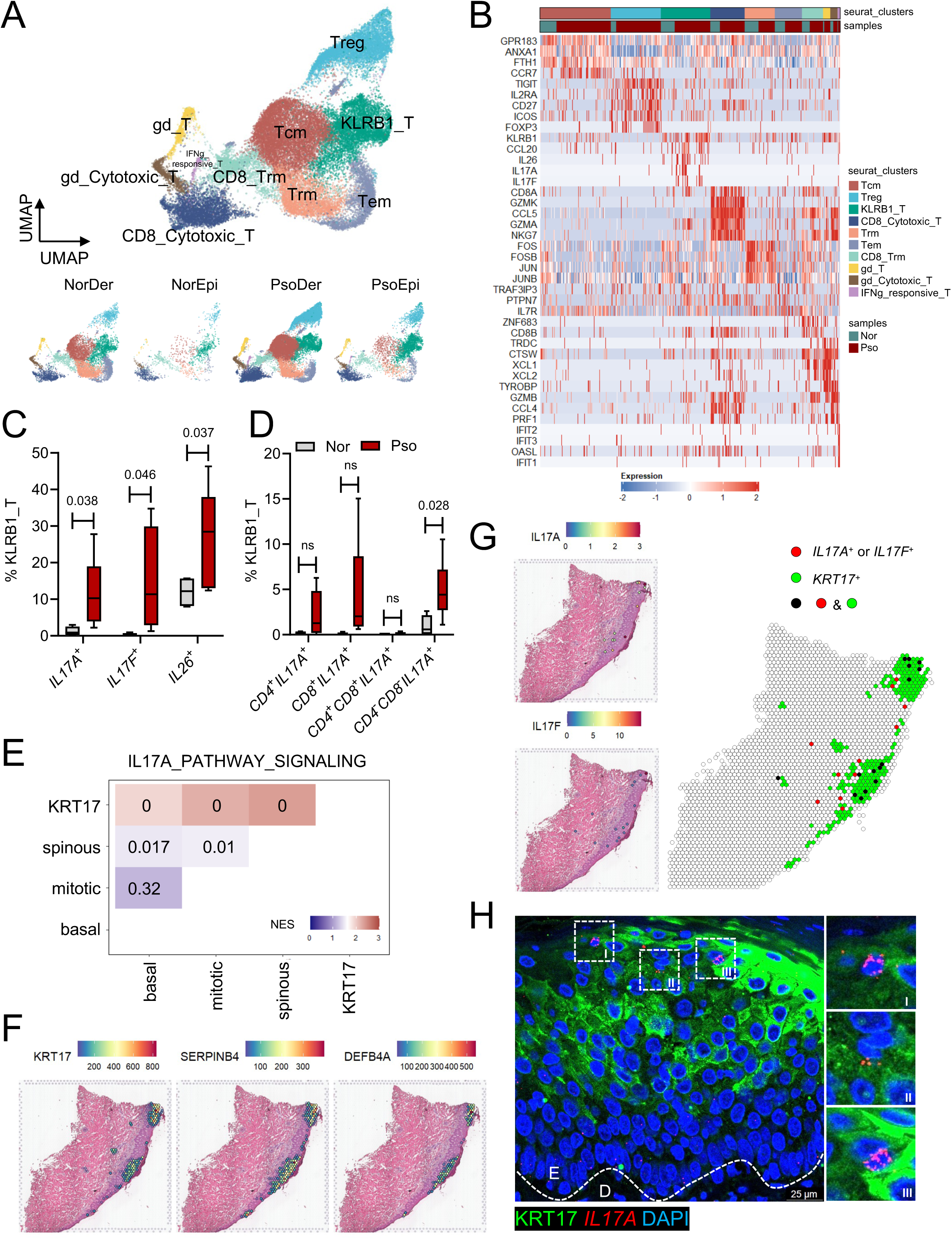
Altered T cell sub-clusters and type 17 biased response in psoriatic skin. (A) UMAP dimensional reduction and sub-clustering of T cells from psoriatic epidermis samples (n = 6), psoriatic dermis samples (n = 6), healthy epidermis samples (n = 4) and healthy dermis samples (n = 4) and split by sample types. (B) Heatmap of signature genes in T cell sub-clusters in (A). (C) Percentages of *IL17A^+^, IL17F^+^* and *IL26^+^* T cells in *KLRB1^+^* T cells in psoriatic samples and healthy controls. Data represent the 25^th^ to 75^th^ percentiles (whiskers showing min to max). *P*-values were determined by two-tailed unpaired *t*-test. ns, not significant. (D) Percentages of *CD4^+^IL17A^+^*, *CD8^+^IL17A^+^*, *CD4^+^CD8^+^IL17A^+^* and *CD4^-^CD8^-^IL17A^+^* T cells in *KLRB1^+^*T cells in psoriatic samples and healthy controls. Data represent the 25^th^ to 75^th^ percentiles (whiskers showing min to max). *P*-values were determined by two-tailed unpaired *t*-test. ns, not significant. (E) GSEA of IL-17A-downstream gene signatures in keratinocyte sub-clusters of psoriatic epidermis. Normalized enrichment score (NES) values were calculated from sub-clusters on the vertical axis versus sub-clusters on the horizontal axis. The numbers represent the nominal (NOM) *P*-values. (F) Spatial feature-plots of indicated genes in psoriatic skin section from a patient. (G) Spatial feature-plots of *IL17A* and *IL17F* in psoriatic skin section from a patient and merged with *KRT17*^+^ (*KRT17* expression > 30) spots. (H) Representative image of RNAscope detection of *IL17A* combined with immunofluorescence of KRT17 in psoriatic skin (n = 3). Scale bar, 25 μm. Data are representative of two independent experiments.

IL-17A induces changes in the expression of psoriasis-associated gene sets in keratinocytes(Muromoto et al., 2016). To dissect the reactivity of keratinocytes to IL-17A, we aggregated the interfollicular keratinocyte sub-clusters as basal (KRT14^+^_ASS1^+^ and KRT14^+^_KRT15^hi^), spinous (KRT10^+^_KRT5^hi^ and KRT10^+^_KRT2^+^), mitotic (PTTG1^hi^ and PCNA^hi^) and KRT17 (KRT17^+^). We then compared these keratinocyte sub-populations using an IL-17A-downstream signature gene set(Muromoto et al., 2016). KRT17 showed enrichment in IL-17A signaling compared to basal, mitotic, and spinous keratinocytes **(Figure 1E**). We also performed spatial transcriptomic analysis on psoriatic skin sections to visualize genes involved in IL-17 responses (**Figure S1A**). In accordance with the gene set enrichment analysis (GSEA), spatial plotting of the IL-17A-downstream signatures showed that *SERPINB4* and *DEFB4A* co-localized with *KRT17* in the outermost layers of psoriatic epidermis **(Figures 1F** and **S4A)**. However, neither spatial plotting nor immunofluorescence staining showed any preferential co-localization of IL17RA with KRT17^+^ keratinocytes **(Figures S4A**-**S4C)**. We hypothesized that in addition to expressing IL17RA, KRT17^+^ keratinocyte induction also required additional IL-17. Importantly, we found that *IL17A^+^* or *IL17F*^+^ spots were intimately co-localized with KRT17^+^ keratinocytes in psoriatic skin (**Figures 1G** and **S5**). *In situ* RNA hybridization of *IL17A* combined with immunofluorescence staining of KRT17 further confirmed the proximity of IL-17-producing T cells with KRT17^+^ keratinocytes (**Figure 1H**). Together, our data suggested that CD161^+^ αβT cell-derived IL-17 acts on adjacent KRT17^+^ keratinocytes to promote and maintain auto-inflammation in psoriasis.

### CCR7^+^ DC are major IL-23 producers in psoriasis

Human DCs and monocytes are highly heterogeneous in their origins and functions(Anderson et al., 2021; Dress et al., 2020), and their contributions to psoriasis pathophysiology remain to be elucidated, although cDC secrete IL-23 as a third signal that induces polarization toward a type 17 T cell response that drives psoriatic plaque formation in mice(Wohn et al., 2013). Based on our transcriptomic analysis, we detected cDC1 (*CLEC9A*^+^*IRF8*^+^), cDC2 (*CLEC10A*^+^*CD1C*^+^), DC3 (*CD14*^+^*CD1C*^+^)(Nakamizo et al., 2021), monocytes/macrophages (*CD14*^+^*C1QA*^+^), CCR7^+^ DC (*CCR7*^+^*LAMP3*^+^*CD200*^+^*IL4I1*^+^)(Maier et al., 2020), LC (*CD1A*^+^*CD207*^+^), Pdc (*JCHAIN*^+^*IRF7*^+^) and AXL^+^ DC (*AXL*^+^)(See et al., 2017) in psoriatic and healthy skin (**Figures 2A** and **S6A**).

**Figure 2.**
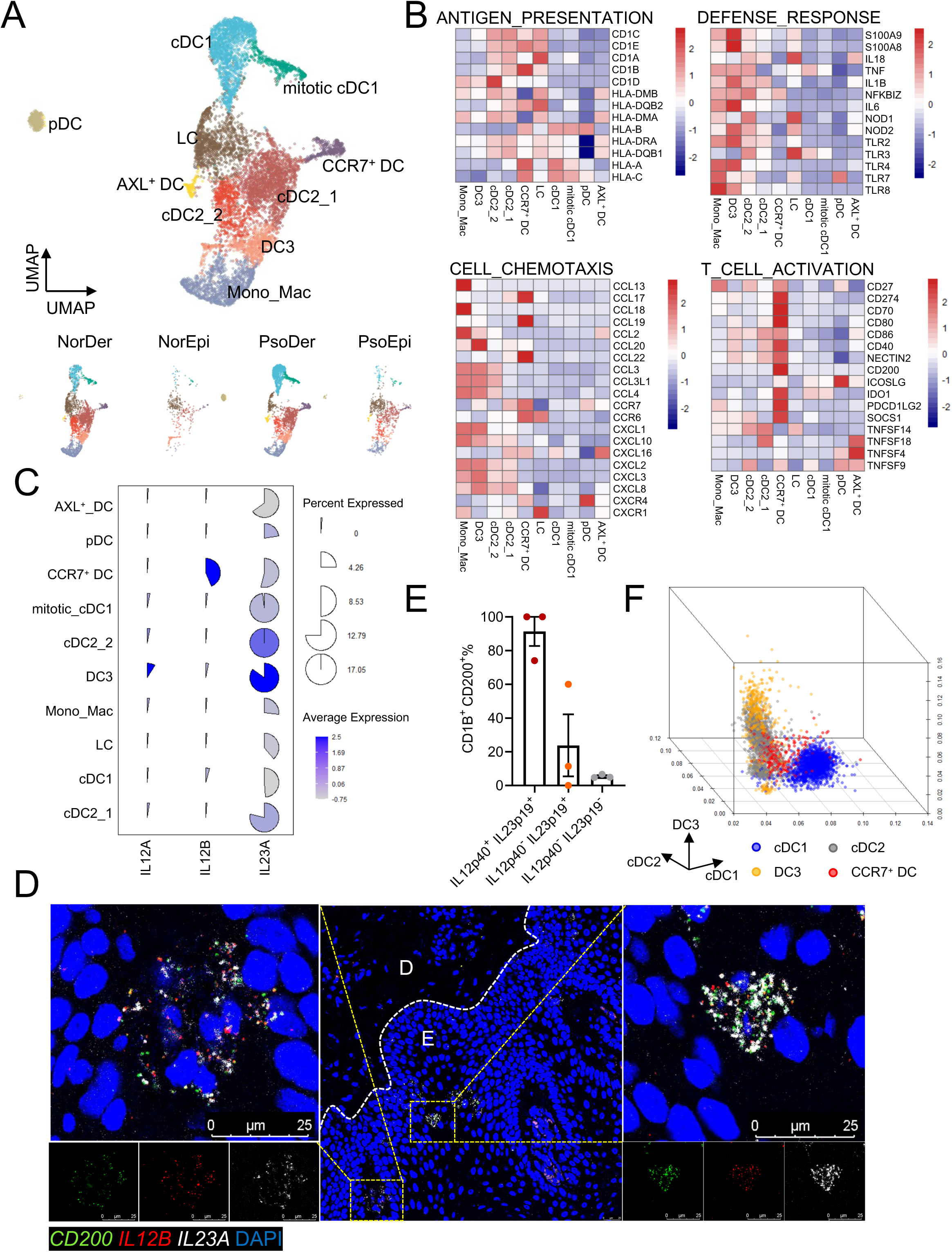
CCR7^+^ DC dominantly produce IL-23 in psoriatic skin. (A) UMAP dimensional reduction and sub-clustering of DCs and monocytes from psoriatic epidermis samples (n = 6), psoriatic dermis samples (n = 6), healthy epidermis samples (n = 4) and healthy dermis samples (n = 4) and split by sample types. (B) Heatmaps of genes associated with the indicated functions in different DC and monocyte subtypes of psoriatic skin. (C) Cell percentages expressing *IL12A*, *IL12B* and *IL23A* and average expression of these genes in different DC and monocyte subtypes of psoriatic skin. (D) Representative image of RNAscope detection of *CD200*, *IL12B* and *Il23A* mRNAs in psoriatic skin (n = 3). Scale bar, 25 μm. Data are representative of three independent experiments. (E) Flow cytometry showing the percentages of CD1B^+^CD200^+^ cells in IL-12p40^+^IL-23p19^+^, IL-12p40^-^IL-23p19^+^ and IL-12p40^-^IL-23p19^-^ cells among CD45^+^CD11C^+^ cells in psoriatic skin samples (n = 3). Data represent the mean ± SEM. Data are representative of three independent experiments. (F) Three-dimensional scatter plots showing cDC1 scores, cDC2 scores and DC3 scores of cDC1, cDC2, DC3 and CCR7^+^ DC in psoriatic skin. The scores were calculated as the fraction of RNA in a cell belonging to genes in the list shown in Supplementary Table 1.

Gene expression analysis of DCs and monocytes in psoriatic skin revealed that CCR7^+^ DC expressed high levels of HLA genes, including *CD1B*, *CD1E*, *HLA-A*, *HLA-B*, and *HLA-C*, and T cell co-stimulatory or co-inhibitory genes, including *CD40*, *CD80*, *CD86*, *CD200*, *CD274*, *CD70* and *PDCD1LG2* (**Figure 2B**). These findings indicated that CCR7^+^ DC express all the key molecules required to activate T cells in psoriatic skin. Moreover, *IL12B* (encoding IL-12p40) and *IL23A* (encoding IL-23p19), which form the intact IL-23 for extracellular secretion(Oppmann et al., 2000), were expressed at high levels by CCR7^+^ DC, but not other types of DCs or monocytes, suggesting that CCR7^+^ DC are potent IL-23-producers and type 17 T cell response-inducers (**Figure 2C**). Single-molecule *in situ* RNA hybridization of *IL23A* and *IL12B* as well as the CCR7^+^ DC marker *CD200* in human psoriatic skin sections revealed a small fraction of *CD200*^+^ cells expressing both *IL23A* and *IL12B* (**Figure 2D**). Flow cytometric analysis of psoriatic skin showed that CD11C^+^ pan DC consisted of IL-12p40^-^IL-23p19^-^, IL-12p40^-^IL-23p19^+^, and IL-12p40^+^IL-23p19^+^ populations (**Figures S6B** and **S6C**), with the IL-12p40^+^IL-23p19^+^ cells confined to the CD1B^+^CD200^+^CCR7^+^ DC population (**Figures S6B** and **2E**). Given that IL-23p19 shows no biological activity without forming a heterodimer with IL-12p40(Oppmann et al., 2000), our data suggested that IL-12p40, but not IL-23p19, is the limiting factor for the production of biologically active IL-23 by DCs in psoriatic skin.

We then studied the lineage of CCR7^+^ DC in psoriatic skin, as these cells were reported to arise from both cDC1 and cDC2, and possibly DC3(Kvedaraite and Ginhoux, 2022). We characterized cDC1, cDC2, DC3 and CCR7^+^ DC based on cDC1, cDC2 and DC3 scores determined according to the following formula: sum [gene values of (cDC1 markers, cDC2 markers, or DC3 markers) in an individual cell]/sum (22,438 annotated gene values in the same cell). Our data suggested that CCR7^+^ DC in psoriatic skin were similar to cDC2 at the transcriptional level (**Figure 2F**). Collectively, our results indicated that CCR7^+^ DC likely arise from cDC2 and provide all the key signals required to activate the T17 cell response in psoriasis.

### CCR7^+^ DC ablation disables IL-23 signaling in psoriasis-like mouse skin

We next used the IL-23-dependent IMQ-induced mouse model of psoriasis to confirm that CCR7^+^ DC is the primary source of competent IL-23(van der Fits et al., 2009). We collected mouse skin samples treated with IMQ for 2 days or 5 days and untreated skin as a control for scRNA-seq. After extraction of the myeloid cells, the integrated data were sub-clustered in an unbiased manner and we named the cells according to their countermarks (**Figures S7A** and **3A**). In accordance with the findings in human psoriatic skin, CCR7^+^ DC, but not other types of DCs or monocytes, expressed both the *Il23a* and *Il12b* transcripts (**Figure 3B**). Our similarity analysis suggested that CCR7^+^ DC resembled cDC2 in both normal and IMQ-treated mouse skin (**Figure 3C**). Previous studies demonstrated that Mgl2^+^ myeloid cell deletion abrogated IL-23 in psoriasis-like skin inflammation(Kim et al., 2018; Whitley et al., 2022), while our data suggested that skin *Mgl2*^+^ myeloid cells consisted of *Ear2*^+^ DC3 and *Apod*^+^ cDC2, with the latter cells possibly giving rise to *Il4i1^+^*CCR7^+^ DC that produced IL-23 (**Figures S7A**, **3C** and **3D**).

**Figure 3.**
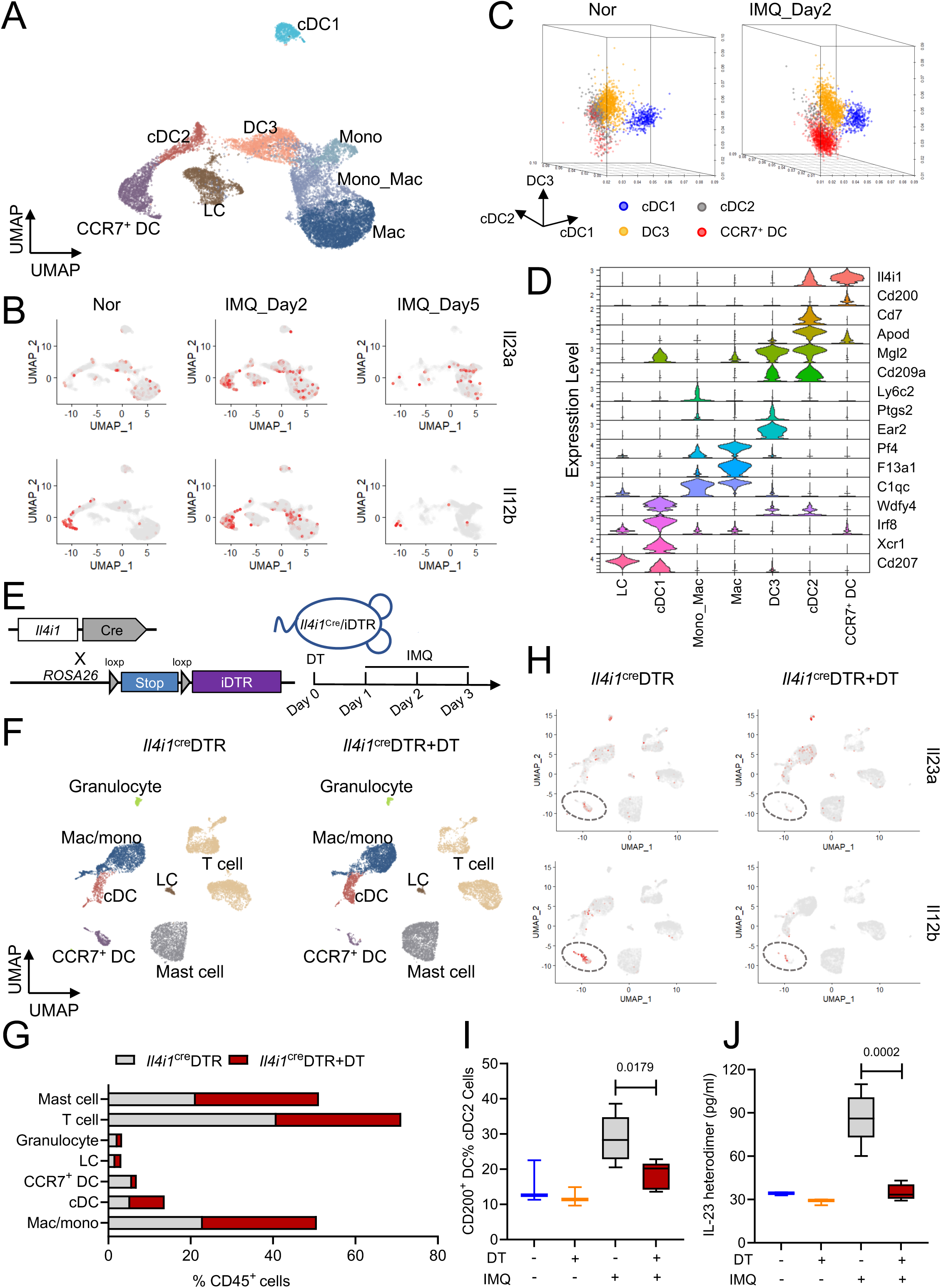
CCR7^+^ DC ablation disables IL-23 signaling in psoriasis-like mouse skin. (A) UMAP dimensional reduction and sub-clustering of scRNA-seq data for sorted CD45^+^Ly6G^-^CD3^-^CD19^-^ cells from mouse ears treated or not with IMQ for 2 days or 5 days. (B) Feature-plots of *Il23a* and *Il12b* expression in (A). (C) Three-dimensional scatter plots showing cDC1 scores, cDC2 scores and DC3 scores of cDC1, cDC2, DC3 and CCR7^+^ DC from mouse ears treated or not with IMQ for 2 days. The scores are calculated as the fraction of RNA in a cell belonging to genes in the list shown in Supplementary Table 1. (D) Violin-plots of indicated genes in different myeloid cells from mouse ears treated with IMQ for 2 days. (E) The strategy to delete CCR7^+^ DC and schematic diagram of DT and IMQ treatment. (F) UMAP dimensional reduction and sub-clustering of scRNA-seq data for ears from IMQ-treated *Il4i1*^cre^DTR mice pre-injected with DT or not. (G) Proportions of different cell types in CD45^+^ cells of ears from IMQ-treated *Il4i1*^cre^DTR mice pre-injected with DT or not. (H) Feature-plots of *Il23a* and *Il12b* expression in (F). (I) Flow cytometry showing the percentages of CCR7^+^ DC in cDC2 (CD45^+^Ly6G^-^CD11c^+^MHCII^+^CD64^-^CD326^-^XCR1^-^) of ears from *Il4i1*^cre^DTR mice treated under the indicated conditions (n = 3∼5). Data represent the 25^th^ to 75^th^ percentiles (whiskers showing min to max). *P*-values were determined by one-way ANOVA. Data are representative of two independent experiments. (J) ELISA quantification of IL-23 in ears from *Il4i1*^cre^DTR mice treated under the indicated conditions (n = 3∼5). Data represent the 25^th^ to 75^th^ percentiles (whiskers showing min to max). *P*-values were determined by one-way ANOVA. Data are representative of two independent experiments.

To study IL-23 in a loss-of-CCR7^+^ DC setting, we generated *Il4i1*^cre^ mice and crossed them with ROSA26iDTR mice (**Figure 3E**). Injection of the *Il4i1*^cre^ DTR mice with diphtheria toxin (DT) resulted in the deletion of IL-23-producing CCR7^+^ DC, as confirmed by both scRNA-seq and flow cytometric analysis (CD45^+^Ly6G^-^CD11c^+^MHCII^+^CD64^-^CD326^-^XCR1^-^CD200^+^) of mouse skin treated with IMQ for 3 days (**Figures 3F**-**3I** and **S7B**). Enzyme-linked immunosorbent assay (ELISA) of the IL-23 heterodimer further confirmed that CCR7^+^ DC deletion abrogated IL-23 expression in IMQ-treated murine skin (**Figure 3J**). Thus, we demonstrated that CCR7^+^ DC are the source of IL-23 in the IMQ-induced mouse model of psoriasis, which is consistent with the scenario in human psoriasis.

### *IL23a* overexpression in Il4i1^+^ cells elicits psoriasis-like skin inflammation

To study CCR7^+^ DC in a gain-of-function setting, we crossed *Il4i1*^cre^ mice with *CAG*-*LSL*-*IL23a* mice and overexpressed IL-23a in CCR7^+^ DC (**Figure 4A**). Importantly, *Il4i1*-*Il23a*^OE^ mice developed scaly plaques on the hairless skin regions, including ears and tails, at 12 weeks of age (**Figure 4B**). Histological examination of the skin lesions showed epidermal hyperplasia (acanthosis) with loss of the granular layer in the epidermis together with accumulation of microabscesses on the surface of the thickened epidermis and massive cellular infiltrates in the dermis (**Figure 4C**). Immunohistochemical analysis of Krt6, Krt5, Krt1/10, Ki67 and filaggrin further confirmed the hyperproliferation and abnormal differentiation of keratinocytes in *Il4i1*-*Il23a*^OE^ mice (**Figure 4D**). We also crossed *Itgax*^cre^ mice with *CAG*-*LSL*-*IL23a* mice to achieve IL-23a overexpression in CD11c^+^ pan DC. The resulting *Itgax*-*Il23a*^OE^ mice developed systemic inflammatory phenotypes, although their psoriasis-like skin inflammatory features, especially acanthosis, were not as prominent as those of the *Il4i1*-*Il23a*^OE^ mice (**Figure S8A**). Thus, we showed that *Il4i1*-*Il23a*^OE^ mice recapitulate key pathological characteristics of human psoriasis in a cell-type-restricted fashion.

**Figure 4.**
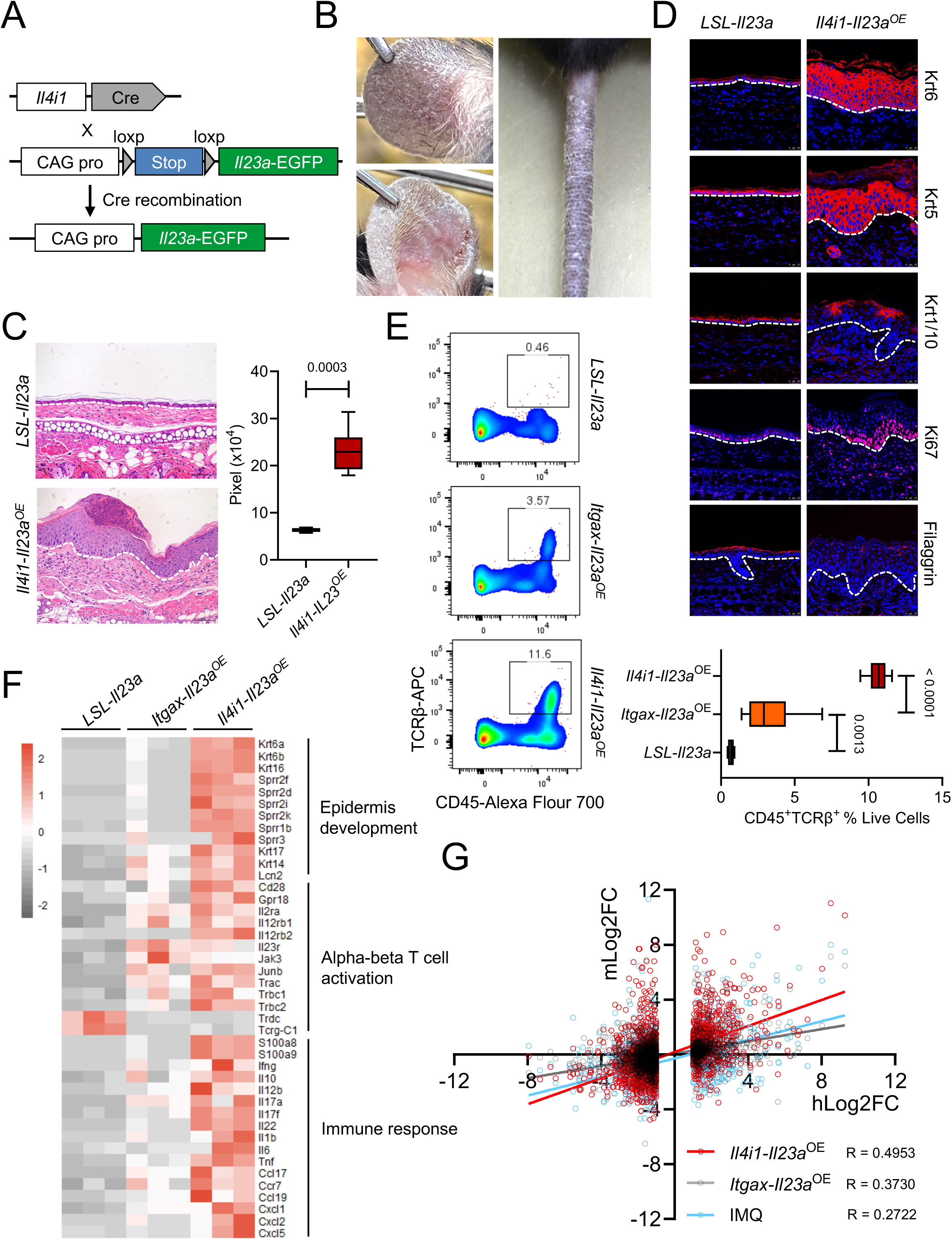
*Il23a* overexpression in CCR7^+^ DC elicits psoriasis-like skin disease. (A) Strategy for developing *Il4i1-Il23a^OE^* mice. (B) Representative macroscopic views of skin lesions of 12-week-old *Il4i1-Il23a^OE^* mice (n = 5). (C) Representative H&E images and quantification of acanthosis in 12-week-old *Il4i1-Il23a^OE^* mice (n = 6) and *LSL-Il23a* mice (n = 5). Scale bar, 50 μm. Data represent the 25^th^ to 75^th^ percentiles (whiskers showing min to max). *P*-values were determined by two-tailed unpaired *t*-test. (D) Representative immunofluorescence images of Krt6, Krt5, Krt1/10, Ki67 and Filaggrin of ears from 12-week-old *LSL-Il23a* mice and *Il4i1-Il23a^OE^* mice (n =3). Scale bar, 25 μm. (E) Flow cytometry showing the percentages of CD45^+^TCRβ^+^ cells in live cells of ears from 12-week-old *LSL-Il23a* mice (n = 8), *Itgax-Il23a*^OE^ mice (n = 6) and *Il4i1-Il23a^OE^* mice (n = 6). Data represent the 25^th^ to 75^th^ percentiles (whiskers showing min to max). *P*-values were determined by one-way ANOVA. (F) Heatmap of selected genes from bulk RNA-seq data from the ears of 12-week-old *LSL-Il23a* mice (n = 3), *Itgax-Il23a*^OE^ mice (n = 3) and *Il4i1-Il23a^OE^*mice (n = 3). The GO categories are indicated. (G) Linear correlations between mouse transcriptional profiles (model versus control) and human transcriptional profile (psoriasis versus healthy control). Genes identified as DEGs (|log2FC| ≥ 1 and *p* < 1×10^−6^) in the human dataset (GSE54456) were used (Li et al., 2014), and mouse genes were joined by case-insensitive gene symbol matching. The gene expression levels of *Il4i1-Il23a^OE^* mice or *Itgax-Il23a*^OE^ mice versus *LSL-Il23a* mice were from our data, and the gene expression levels of IMQ-treated C57BL/6J mice versus control mice were from GSE86315 (Swindell et al., 2017). Data (**B**-**E**) are representative of three independent experiments.

Psoriasis mouse models, including IMQ-induced and recombinant IL-23-mediated skin inflammation, differ from human psoriasis in that IL-17 is produced mainly by γδT cells rather than αβT cells in these models(Cai et al., 2011; Mabuchi et al., 2011). Notably, *Il4i1*-*Il23a*^OE^ mouse skin contained significantly more αβT cells than *Itgax*-*Il23a*^OE^ mouse skin (**Figure 4E**), and IL-17a^+^ T cells (T17) were mostly αβT cells in the skin lesions of *Il4i1*-*Il23a*^OE^ mice (**Figure S8B**). Moreover, transcriptomic profiling of *LSL*-*Il23a*, *Itgax*-*Il23a*^OE^ and *Il4i1*-*Il23a*^OE^ mouse skin revealed human psoriasis-like gene alterations in *Il4i1*-*Il23a*^OE^ mice, with the differentially expressed genes (DEGs) enriched in epidermis development (*Krt6a*, *Krt6b*, *Krt16* and *Krt17*), αβT cell activation (*Trac*, *Trbc1*, *Trbc2*, *Cd28* and *Gpr18*), and immune response (*Il12b*, *Il17a*, *Il17f* and *Il22*, **Figure 4F**). The DEGs in *Il4i1*-*Il23a*^OE^ mouse skin resembled those in human psoriatic skin when analyzed using a linear regression model, which showed a stronger coefficient of determination (R) compared to those in IMQ-treated or *Itgax*-*Il23a*^OE^ mouse skin (**Figure 4G**). Collectively, our findings demonstrated that IL-23a overexpression in Il4i1^+^ cells leads to a psoriasis-like mouse model characterized by αβT cell activation, thus highlighting the central role of CCR7^+^ DC in the cellular and molecular program that drives psoriasis.

### *Il4i1*-*Il23a*^OE^ mice develop psoriatic arthritis-like symptoms

Up to 30% of psoriasis patients develop psoriatic arthritis (PsA), which is diagnosed according to inflammatory musculoskeletal features in the joints, entheses or spine in the presence of skin and/or nail psoriasis(FitzGerald et al., 2021). We observed digit swelling (dactylitis) and paw swelling in *Il4i1*-*Il23a*^OE^ mice after 12 weeks of age, and histological examination of the paws confirmed transformation of the synovial lining into hyperplastic pannus in the joints (**Figures 5A** and **5B**). Bulk RNA-seq of the metacarpal and phalangeal bones of the fore paws confirmed the presence of bone resorption, as revealed by the increased expression of *Oscar, Fcgr4*, *Dcstamp* and *Adam8*. Furthermore, key drivers of bone resorption including *Tnfsf11* (RANKL), *Ocstamp*, *Tyobp*, *Itgb3*, *Slc9b2* were significantly enriched in the bones and joints of *Il4i1*-*Il23a*^OE^ mice, indicating that this effect was due to osteoclast activation (**Figure 5C**). The data further revealed neutrophil infiltration and IFN-γ-induced signaling in the bones and joints of *Il4i1*-*Il23a*^OE^ mice, suggesting a resemblance to the immunophenotype involving mixed type 1 and type 17 T cell responses in human PsA (**Figure 5C**)(FitzGerald et al., 2021). We then visualized the structural damages of hind paws from 25∼29-week-old *Il4i1*-*Il23a*^OE^ as compared to *LSL*-*Il23a* control mice using microCT (**Figure 5D**). Scanning of the intersecting surfaces of the metatarsal bones showed decreased cortical bone thickness (Ct.Th), cortical bone area (Ct.Ar) and bone volume to total volume ratio (BV/TV) in *Il4i1*-*Il23a*^OE^ mice (**Figures 5E** and **5F**). These data clearly demonstrated bone destruction of the metatarsophalangeal joint, which is the most affected site in PsA patients(Wang et al., 2023). Thus, *Il4i1*-*Il23a*^OE^ mice were validated as a model of PsA, with pathological features including T cell activation, osteoclast differentiation, and bone resorption in the bones and joints.

**Figure 5.**
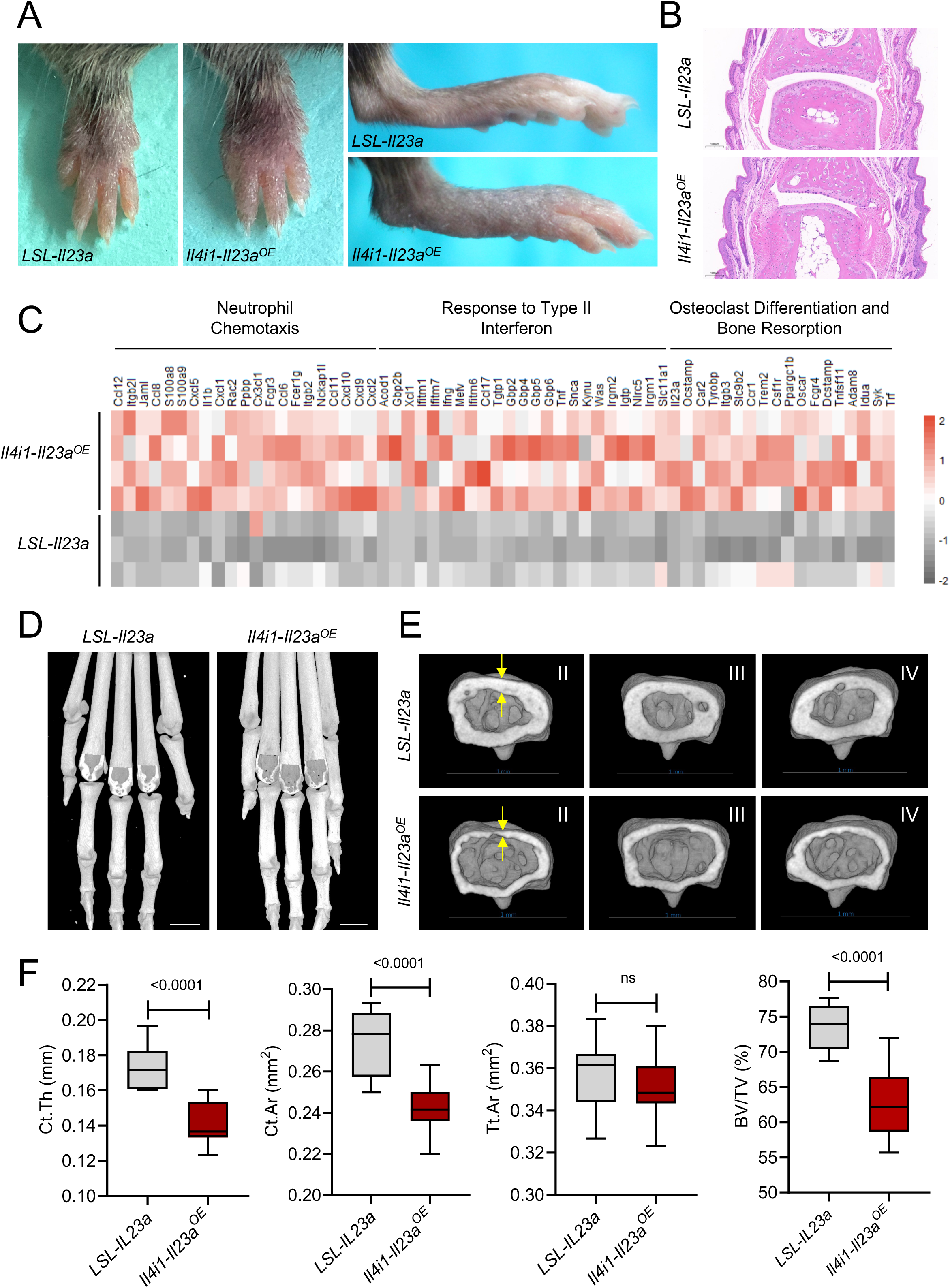
*Il4i1*-*Il23a*^OE^ mice develop psoriatic arthritis-like symptoms. (A) Representative macroscopic views of the paws of 16-week-old *LSL-Il23a* mice (n = 5) and *Il4i1-Il23a^OE^* mice (n = 5). (B) Representative H&E images of the hind paws from 16-week-old *LSL-Il23a* mice (n = 3) and *Il4i1-Il23a^OE^* mice (n = 6). Scale bar, 100 μm. (C) Heatmap of selected genes from bulk RNA-seq data of the metacarpal and phalangeal bones from 16-week-old *LSL-Il23a* mice (n = 3) and *Il4i1-Il23a^OE^*mice (n = 4). The GO categories are indicated. (D) Representative microCT images of the hind paws from 25∼29-week-old *LSL-Il23a* mice (n = 8) and *Il4i1-Il23a^OE^* mice (n = 11). Scale bar, 1 mm. (E) Representative microCT images of the metatarsal bones proximal to the metatarsophalangeal joints from 25∼29-week-old *LSL-Il23a* mice (n = 8) and *Il4i1-Il23a^OE^* mice (n = 11). II, III and IV represent three different toes of the mice. Scale bar, 1 mm. (F) Quantification of bone structural parameters of 25∼29-week-old *LSL-Il23a* mice (n = 8) and *Il4i1-Il23a^OE^* mice (n = 11) shown in (E). Data represent the 25^th^ to 75^th^ percentiles (whiskers showing min to max). Ct.Th, cortical bone thickness; Ct.Ar, cortical bone area; Tt.Ar, total area of the cross section; BV/TV, bone volume to total volume ratio; ns, not significant. Data (**A**, **B** and **D**-**F**) are representative of three independent experiments.

### Spatial crosstalk between CCR7^+^ DC, CD161^+^ T cells and KRT17^+^ keratinocytes in psoriasis

Having demonstrated that CD161^+^ T cell-derived IL-17 regulated KRT17^+^ keratinocytes, and CCR7^+^ DC-derived IL-23 is a prerequisite of IL-17 production, we next explored the relationships between these three cell types and visualized CD161^+^ T cells (*KLRB1*^+^) and CCR7^+^ DC (*LAMP3*^+^) in psoriatic skin sections. Importantly, in psoriatic epidermis, CCR7^+^ DC were co-localized with CD161^+^ T cells (**Figures 6A** and **S9**). Immunofluorescence staining of LAMP3, CD161 and KRT17 confirmed the spatial transcriptome data (**Figure 6B**), in that CCR7^+^ DC, CD161^+^ T cells, and KRT17^+^ keratinocytes form a spatial cellular module that sustains the IL-23-T17 inflammatory axis in psoriasis. To further clarify the mechanism underlying the formation of such a module, we studied cell chemotaxis in psoriatic skin by predicting cell-cell ligand-receptor interactions using CellPhoneDB(Efremova et al., 2020). Importantly, epidermal CCR7^+^ DC expressed *CCL19*, which is required to recruit dermal CCR7^+^ DC, and *CXCL16*, which attracts dermal and epidermal CD161^+^ T cells (**Figures 6C** and **6D**). Taken together, our data supported a working model in which CCR7^+^ DC first reach the epidermis to initiate the IL-23-dominated inflammatory program that involves the subsequent recruitment of CD161^+^ T cells and the production of IL-17.

**Figure 6.**
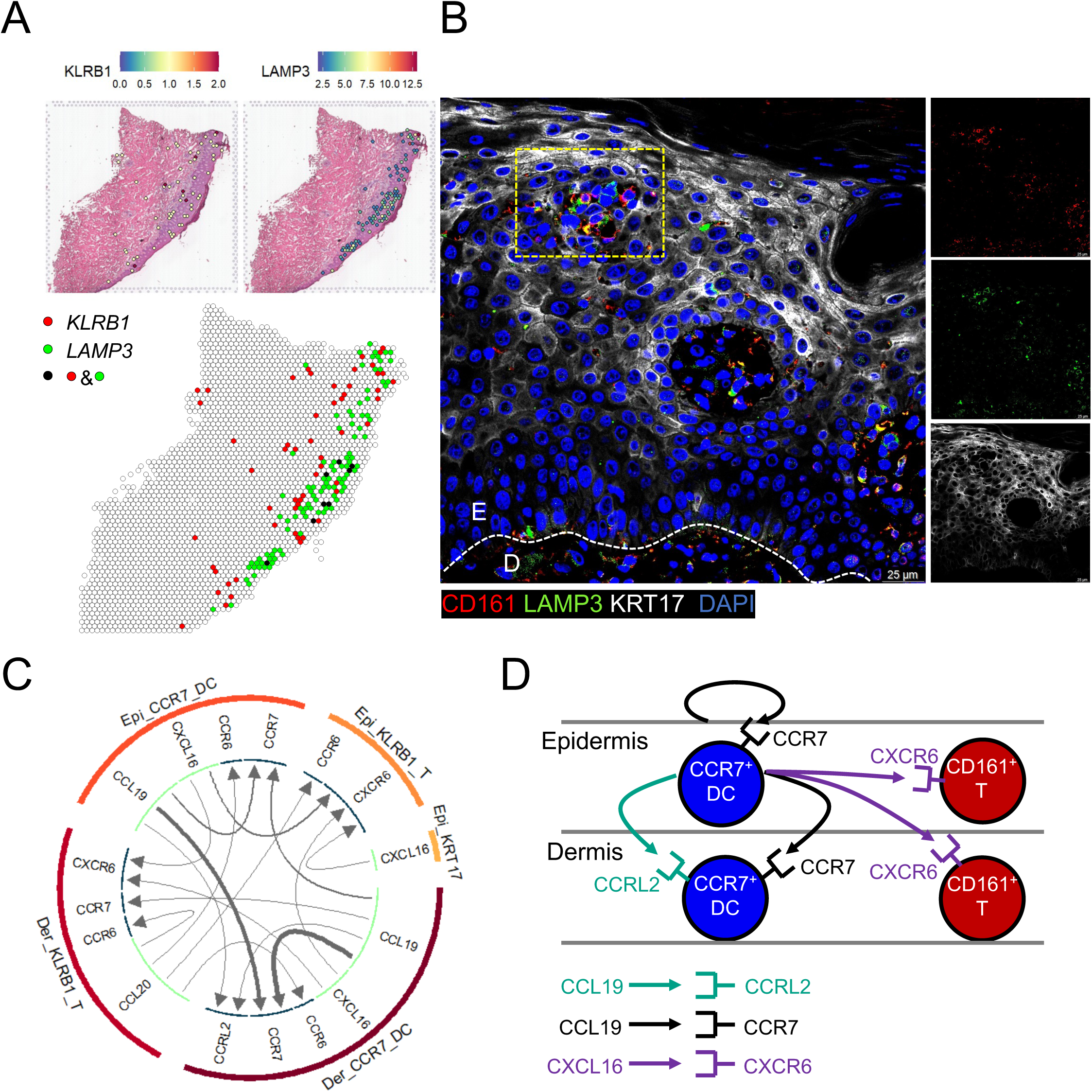
CCR7^+^ DC define a type 17 spatial module in psoriatic epidermis. (A) Spatial feature-plots of *KLRB1* and *LAMP3* in psoriatic skin section from a patient. (B) Representative immunofluorescent labeling of CD161, LAMP3 and KRT17 in psoriatic skin (n = 3). Data are representative of two independent experiments. (C) Chemokine receptor-ligand pairs across cell subpopulations within psoriatic skin. All interactions shown are statistically significant (*p* < 0.05), and arrows denote directionality from ligand to receptor. (D) A working model of chemotaxis of CCR7^+^ DC and CD161^+^ T cells in psoriatic skin.

## DISCUSSION

In this study, we reveal the immune landscapes of the epidermis and dermis in psoriasis compared with homeostasis. We show that the hyperproliferative keratinocytes in psoriasis, which undergo a disordered differentiation program, are characterized by ectopic overexpression of KRT17 in their most differentiated state. Localized in the outermost layers of psoriatic epidermis, KRT17^+^ keratinocytes express high levels of IL-17-downstream gene signatures, which is attributed to their spatial proximity to IL-17-producing CD161^+^ T cells. In psoriatic skin, CCR7^+^ DC, which are likely differentiated from cDC2, enter the epidermis and recruit CD161^+^ T cells. Importantly, CCR7^+^ DC readily provide first, second and third signals (IL-23) that activate CD161^+^ T cells and induce the production of IL-17. The intimate co-localization of CCR7^+^ DC, IL-17-producing CD161^+^ T cells, and KRT17^+^ keratinocytes therefore indicates a type 17 spatial module in psoriatic epidermis, suggesting pathogenic roles of cells within the module, as well as the therapeutic potential of harnessing this module in psoriasis.

The cellular source of IL-23 in psoriatic skin is controversial. Several studies showed that monocytes/macrophages or monocyte-derived DCs express IL-23A(Cai et al., 2011; Fuentes-Duculan et al., 2010; Hansel et al., 2011), while more recently, Nakamizo et al. reported that CD14^+^ DC3 express a high level of IL-23A in human psoriatic skin(Nakamizo et al., 2021). Studies on mouse models of psoriasis suggested that Mgl2 (CD301b)^+^ cDC2 were the major source of IL-23(Kim et al., 2018; Whitley et al., 2022); however, there is no human homolog of the *Mgl2* gene. Due to the enigma related to the source of IL-23, the upstream regulatory mechanisms of IL-23 production are ill-defined, thus hindering the development of targeted therapies focusing on modulating IL-23 expression. In this study, we showed that IL-23A is indeed expressed by many monocyte and DC populations with CD14^+^ DC3 harboring the maximum transcripts, while IL-12B is dominantly expressed by CCR7^+^ DC in human psoriatic skin. Given that IL-23A cannot be secreted and thus, is not biologically active without binding to IL-12B(Oppmann et al., 2000), our findings demonstrate that the cellular specificity of IL-23 production is determined by IL-12B rather than IL-23A, and that CCR7^+^ DC are the main producer of IL-23 in psoriasis.

Here, we employed scRNA-seq to confirm that CCR7^+^ DC predominantly produce IL-23 in the IMQ-induced mouse model of psoriasis. Notably, *Mgl2*^+^ DC consist of *Ear2*^+^ DC3 and *Apod*^+^ cDC2, and the latter possibly differentiate into CCR7^+^ DC in psoriasis-like skin lesions. Because CCR7^+^ DC specifically express high levels of *Il4i1*, we generated *Il4i1*^cre^ mice and crossed them with both ROSA26iDTR and *LSL*-*IL23a* mouse strains to study IL-23 production in both loss and gain-of-function settings. DT-induced deletion of CCR7^+^ DC in *Il4i1*^cre^ DTR mice abrogated IMQ-induced IL-23 expression, implicating CCR7^+^ DC as the main source of IL-23 in psoriasis-like mouse skin. On the other hand, IL-23a overexpression in Il4i1^+^ cells recapitulates key features of human psoriasis including αβT cell-dominated IL-17 production and the propensity to develop PsA. The observation that IL-23a overexpression in CD11c^+^ pan DC does not drive psoriasis-like skin inflammation reveals not only a unique pathogenic role of CCR7^+^ DC, but also offers a valuable mouse model suitable for the investigation of IL-23 generation mechanisms. In addition, this model provides opportunities for the design of new therapies targeting IL-23-producing cells to treat a broad spectrum of auto-inflammatory diseases. Thus, our model represents the first step in developing the next generation of targeted drugs for the purpose of achieving longer term disease remission.

Other than producing IL-23, psoriatic CCR7^+^ DC express high levels of antigen presentation molecules, including *HLA-A*, *HLA-B*, *HLA-C*, *CD1B*, and *CD1E*, and co-stimulatory signals, including *CD80*, *CD86*, and *CD40*. These data suggest the strong antigen-presenting capacity of CCR7^+^ DC and their central role in maintaining the T17 cell response, which is further evidenced by the co-localization of CCR7^+^ DC, CD161^+^ T cells and KRT1*7*^+^ keratinocytes. This “type 17 module” concept and the fact that psoriasis is strongly associated with HLA polymorphisms highlight the importance of further investigation of antigens presented by CCR7^+^ DC and then recognized by CD161^+^ T cells, which may elucidate the exact cause of auto-inflammation in psoriasis and the mechanisms of relapse.

## ACKNOWLEDGMENTS

This work was supported by the National Natural Science Foundation of China Original Exploration Program (82050009), the National Key Research and Development Program of the Ministry of Science and Technology (2020YFA0112900), the National Science Foundation of China (81930088, 82173417 [Y.S.], 82203914 [F.L.] and 82373470 [F.L.]), Shanghai Scientific and Technological Innovation Action Plan (22140903100, 22QA1407600 [F.L.] and 23ZR1480700 [F.L.]), SJTU Trans-med Awards Research (20210102), and Innovative Research Team of High-Level Local Universities in Shanghai [by H.W. if not otherwise noted].

## AUTHOR CONTRIBUTIONS

Conceptualization, H.W., Y.S. and F.L.; Investigation, Y.S., F.L., X.C., Z.W., X.Y., L.S. and Z.L.; Data curation, Y.S.; Writing – original draft, F.L.; Writing – review & editing, H.W., F.G., F.L. and Y.S.; Funding acquisition, H.W., Y.S. and F.L.; Resources, Z.W. and Y-L.S.; Supervision, H.W.; Project administration, H.W.

## EXPERIMENTAL MODEL AND SUBJECT DETAILS

### Human subjects

Psoriatic skin samples were obtained by punch biopsy from patients under local lidocaine anesthesia. Normal adult human skin specimens were obtained from healthy donors undergoing plastic surgery. All participants provided written informed consent. This study was performed in accordance with the principles of the Declaration of Helsinki and approved by the Research Ethics Boards of Shanghai General Hospital, China (No. 2018KY239).

### Animals

C57BL/6 mice were purchased from Shanghai SLAC Laboratory Animal Co., Ltd. *Il4i1*-*2A*-*Cre* (*Il4i1*^cre^) mice and *R26*-*CAG*-*LSL*-*Il23a*-*IRES*-*EGFP* (*LSL*-*Il23a*) mice were produced by Shanghai Model Organisms Co., Ltd. *ROSA26-LSL-DTR* (ROSA26iDTR) mice and *Itgax*^cre^ mice were obtained from Cyagen Co., Ltd. The mice were bred and maintained under specific pathogen-free (SPF) conditions. Age-matched and sex-matched mice were used for all the experiments in accordance with the National Institutes of Health Guide for the Care and Use of Laboratory Animals with the approval (SYXK-2019-0028) of the Scientific Investigation Board of Shanghai General Hospital. Throughout these experimental studies, all efforts were made to alleviate any suffering and the mice were euthanized by CO_2_ inhalation.

## METHOD DETAILS

### Single-cell transcriptomics

#### Sample preparation, library production and RNA sequencing

Fresh skin biopsies were placed in saline at 4L prior to processing. The epidermis was separated from the dermis by dispase II digestion overnight at 4L. Single-cell suspensions were generated by enzyme digestion as previously reported(Lou et al., 2020) and analyzed by flow cytometry to exclude doublets, debris, and DAPI-positive dead cells. For dermis samples, CD45^+^ cells were sorted for subsequent processing. Sorted cells were centrifuged and resuspended in 0.04% BSA in phosphate-buffered saline (PBS). Chromium Single Cell 3’ v3 (10× Genomics) libraries were prepared using the Chromium Controller according to the manufacturer’s instructions. The resulting libraries were sequenced with the Illumina NovaSeq 6000 platform. Trimmed data were processed using the CellRanger (version 3.0, 10× Genomics) and further filtered, processed, and analyzed using the Seurat package (version 4.3.0)(Butler et al., 2018).

#### Data processing and clustering with Seurat

Cells with fewer than 200 genes, more than 5,000 genes, or more than 5% mitochondria content were removed. Doublets were predicted using DoubletFinder and removed(McGinnis et al., 2019). The filtered data were normalized using a scaling factor of 10,000 to generate transcripts per kilobase million (TPM)-like values. We integrated the filtered samples using the FindIntegrationAnchors and IntegrateData functions with default parameters (dimensionality = 30). The top 2,000 most variable genes were selected using the FindVariableFeatures function and the genes were then used for principal component analysis (PCA). The number of PCs for clustering was selected based on the ‘Elbow plot’ of different datasets. Clustering was performed using the FindClusters function with a resolution selected for different datasets. Results were visualized using the Seurat package.

#### Pseudo-time trajectory analysis using Monocle

Keratinocytes pre-clustered and labeled according to the countermark genes were used as an input for Monocle 2 in the pseudo-time trajectory analysis(Trapnell et al., 2014). Genes with expression levels lower than 0.1 and genes expressed by fewer than 10 cells were removed. The remaining cells were clustered in an unsupervised manner and DEGs were identified using the differentialGeneTest function. The top 1,000 significant DEGs were selected as the ordering genes. The DDRTree algorithm was used for dimension reduction and BEAM was used to identify the genes driving the transition in pseudo-time.

### Spatial transcriptomics

Fresh skin biopsies from healthy donors and patients with psoriasis were embedded in optical cutting tissue (OCT) compound and snap-frozen on dry ice. Skin sections (10 μm thick) were prepared using a cryostat microtome and mounted onto Visium slides (Visium Spatial Tissue Optimization Slide & Reagent kit, 10× Genomics). After hematoxylin and eosin (H&E) staining, bright-field images were obtained. Optimized permeabilization (for 24 min) and tissue removal were conducted on the Visium slides. After reverse transcription, the barcoded cDNA was enzymatically released and collected. The cDNA libraries were then sequenced on the Illumina NovaSeq 6000 platform. The data were processed with the SpaceRanger (version 1.1.0, 10× Genomics) and mapped to the GRCh38-2020-A genome. Results were visualized using the Seurat package (version 4.3.0).

### Flow cytometry

Single cell suspensions were generated from the skin of patients with psoriasis, *Il4i1*^cre^ DTR mice, *LSL*-*Il23a* mice, *Itgax*-*Il23a*^OE^ mice and *Il4i1*-*Il23a*^OE^ mice as previously reported(Lou et al., 2020). Cells were stained with fluorophore-conjugated antibodies and assayed with a BD LSRFortessa™ or a BD FACSymphony™ A3 cytometer, and the data were analyzed using FlowJo software. For human sample analyses, antibodies against CD45 (clone 2D1) and CD1b (clone SN13) were obtained from BioLegend; CD45 (clone HI30), CD200 (clone MRC OX-104) and CD11C (clone B-ly6) were obtained from BD Biosciences; IL-23p19 (clone 23dcdp), IL-12/IL-23p40 (clone eBioHP40) were obtained from eBioscience. For mouse sample analyses, antibodies against CD45 (clone 30-F11) were obtained from BioLegend or BD Biosciences; Ly-6G (clone 1A8) and CD200 (clone OX-90) were obtained from BD Biosciences; CD11c (clone N418), TCRbeta (clone H57-597) and MHC Class II (I-A/I-E) (clone M5/114.15.2) were obtained from eBioscience; CD326 (Ep-CAM) (clone G8.8), XCR1 (clone ZET) and CD64 (FcγRI) (clone X54-5/7.1) were obtained from BioLegend.

### RNAscope™ Multiplex Fluorescent Assay

Human paraffin sections were dewaxed, and single-molecule fluorescence *in situ* hybridization (FISH) experiments were carried out using an RNAscope™ Multiplex Fluorescent Assay v2 with an RNAscope™ Probe-Hs-IL23A-C3 (ACD cat. 562851-C3), an RNAscope™ Probe-Hs-IL12B-C2 (ACD cat. 402071-C2), an RNAscope™ Probe-Hs-CD200 (ACD cat. 410471), and an RNAscope™ Probe-Hs-IL17A (ACD cat. 310931). The sections were mounted with a fluorescent mounting medium (Sigma-Aldrich cat. DUO82040) and visualized under a confocal microscope (Leica, STELLARIS 8 DIVE).

### Mouse models of psoriasis

For the IMQ-induced mouse model of psoriasis, male C57BL/6 mice (aged 7 weeks) were maintained under SPF conditions. The mice received a daily topical dose of 25Lmg IMQ cream (5%) (MedShine cat. 120503) per ear for two or five consecutive days before the mice were euthanized and the ears were collected for scRNA-seq. For the *Il4i1*^cre^ DTR mice, 100 μg DT (Sigma-Aldrich cat. D0564) was injected intraperitoneally (i.p.) one day before 25Lmg IMQ cream was applied per ear for two days. The mice were then euthanized and the ears were collected for scRNA-seq, flow cytometry and ELISA. After euthanization, ears were collected from *LSL*-*Il23a*, *Itgax*-*Il23a*^OE^, and *Il4i1*-*Il23a*^OE^ mice (aged 12 weeks) for histological analysis, flow cytometry and bulk RNA-seq, while paws were collected from mice (aged 16 weeks) for histological analysis, microCT and bulk RNA-seq.

### ELISA

Ears from *Il4i1*^cre^ DTR mice were snap-frozen, pulverized, and homogenized for protein extraction using a ProteinExt® Mammalian Total Protein Extraction Kit (Transgen cat. DE101-01). IL-23 heterodimer levels were measured using a Mouse IL-23 Quantikine ELISA Kit (R&D Systems cat. M2300) according to the manufacturer’s instructions.

### Histological analysis and immunofluorescence

Ears from *LSL*-*Il23a*, *Itagx-Il23a*^OE^ and *Il4i1*-*Il23a*^OE^ mice were embedded in paraffin and sectioned (5 μm thick). The sections were stained with H&E, and the pixel size of the epidermal area was measured using the lasso tool in Adobe Photoshop CS4. For immunofluorescence staining, the sections were deparaffinized and retrieval was performed by heating the sections in sodium citrate buffer (pH = 6.0) or Tris-EDTA buffer (pH = 8.0). The sections were blocked for 1 h at room temperature (RT) in PBS containing 1% bovine serum albumin (BSA), 5% goat serum, 0.3% Triton X-100 and stained overnight at 37L in blocking buffer containing primary antibody (anti-Krt6, Polyclonal, Proteintech cat. 10590-1-AP, 1:200 dilution; anti-Krt5, Clone 2C2, Invitrogen cat. MA5-17057, 1:200 dilution; anti-Krt1/10, Clone LH1, Santa Cruz cat. sc-53251, 1:200 dilution; anti-Ki67, Polyclonal, Servicebio cat. GB111499, 1:500 dilution; anti-Filaggrin, Clone FLG01, GeneTex cat. GTX23137, 1:100 dilution; anti-CD161, Clone 14F1F11, NOVUS Cat. NBP2-14845; anti-DC-LAMP, Colne 1010E1.01, NOVUS Cat, DDX0191P-100, anti-KRT17/CK17/Cytoketatin 17, Clone E3, LS Bio Cat. LS-B7169; anti-KRT17/CK17/ Cytoketatin 17, Polyclonal, LS Bio Cat#LS-B7610). Thereafter, the sections were rinsed three times in PBS and stained with fluorochrome-conjugated secondary antibodies (all from Life Technologies, 1:1,000 dilution) for 1 h at RT in the dark. Sections were also stained with DAPI (BD Biosciences cat. 564907, 1:2,000 dilution) at RT for 5 min to visualize nuclei. The sections were mounted with a fluorescent mounting medium (Sigma-Aldrich cat. DUO82040) and visualized under a confocal microscope (Leica, STELLARIS 8 DIVE).

### Bulk RNA-seq

Ears or peeled metacarpal and phalangeal bones were snap-frozen in liquid nitrogen and pulverized. Total RNA was isolated using RNAiso Reagent (TaKaRa cat. 9108). cDNA libraries were prepared using a VAHTS Universal V8 RNA-seq Library Prep Kit for Illumina (Vazyme cat. NR605-01) according to the manufacturer’s instructions and sequenced on a NovaSeq 6000 (Illumina). The adaptor sequences were trimmed from the raw paired-end reads using Skewer(Jiang et al., 2014). The sequences were then aligned to GRCm38 using STAR(Dobin et al., 2013) and assembled using StringTie(Pertea et al., 2015). Gene enrichment analysis (GSEA) was performed to identify enriched pathways(Mootha et al., 2003; Subramanian et al., 2005).

### MicroCT

Mouse hind paws were collected and fixed in 4% paraformaldehyde and analyzed by microCT (Venus001, Pingseng Scientific). Bone structural parameters including total area (Tt.Ar), cortical bone thickness (Ct.Th), cortical bone area (Ct.ar) and cortical bone area to total cross-sectional area ratio (Ct.ar/Tt.ar) were calculated using DataViewer software (Bruker). The samples were then decalcified using 10% EDTA and embedded for H&E staining.

## QUANTIFICATION AND STATISTICAL ANALYSIS

Data were presented as the 25^th^ to 75^th^ percentiles (whiskers showing min to max) or as the mean ± SEM and analyzed using GraphPad Prism 9. Differences between two groups were evaluated using Student’s *t*-test, and differences between multiple groups were evaluated by analysis of variance (ANOVA) with Geisser-Greenhouse correction. A simple linear regression model was used to analyze the transcriptome correlation between mouse models and human psoriasis.

## DATA AVAILABILITY

The sequencing data in this paper are deposited in Genome Sequence Archive (GSA). The bulk RNA sequencing data are deposited under the accession: CRA013603; the human single-cell transcriptomics data are deposited under the accessions: HRA003418 and HRA006130; the human spatial transcriptomics data are deposited under the accession: HRA006129.

---

**Figure S1.**
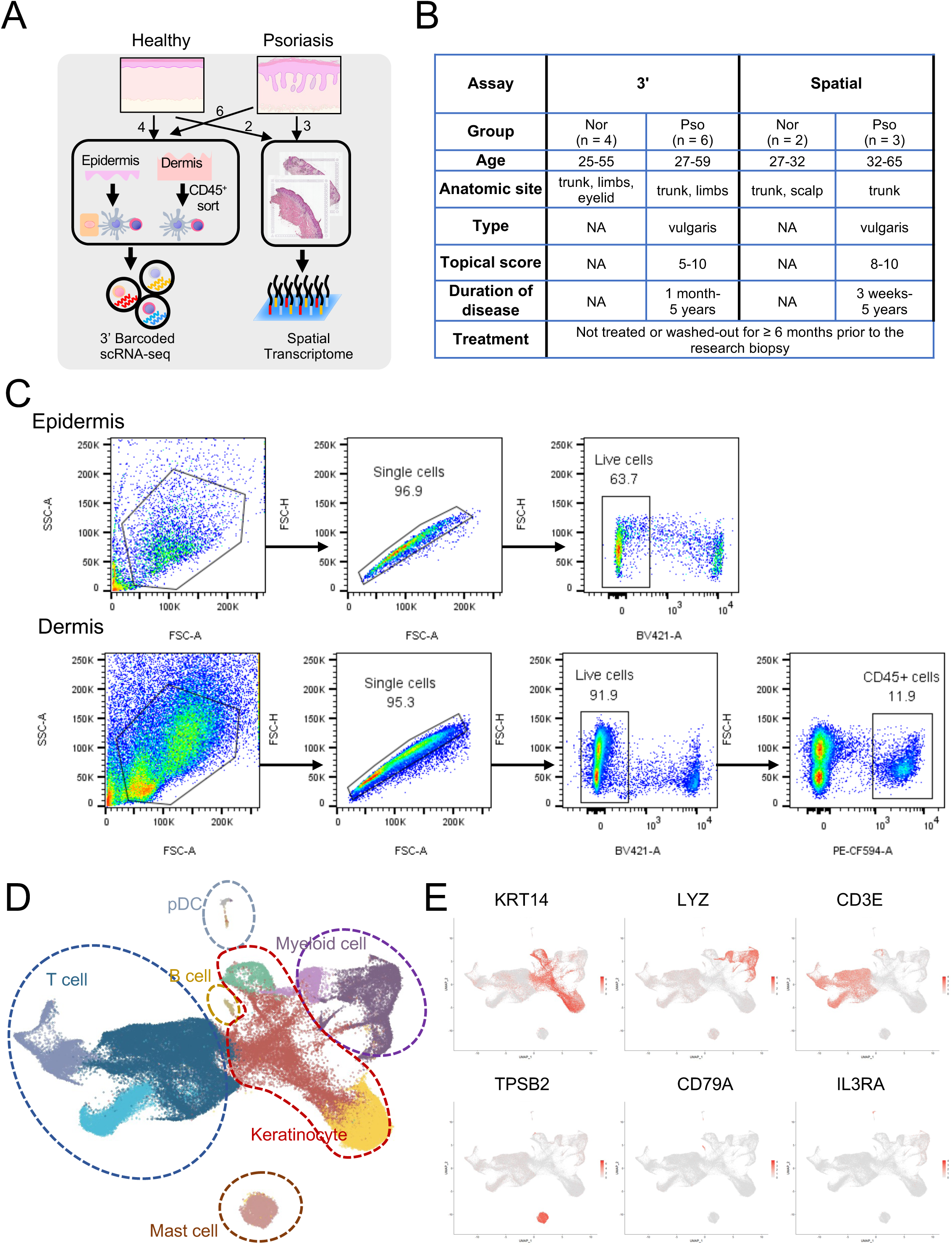
A single-cell transcriptomic landscape in the skin of patients with psoriasis and healthy donors. (A) Schematic overview of skin samples processed for 3’-barcoded scRNA-seq and spatial transcriptome. (B) General information of patients with psoriasis and healthy donors. (C) Representative flow cytometry plots showing the gating strategy of all live cells in the epidermis and CD45^+^ live leukocytes in the dermis. (D) UMAP dimensional reduction and cell type clustering of 3’-barcoded scRNA-seq data containing epidermal cells and dermal CD45^+^ leukocytes from patients with psoriasis and healthy donors. (E) Feature-plots of marker genes for cell clustering in (D).

**Figure S2.**
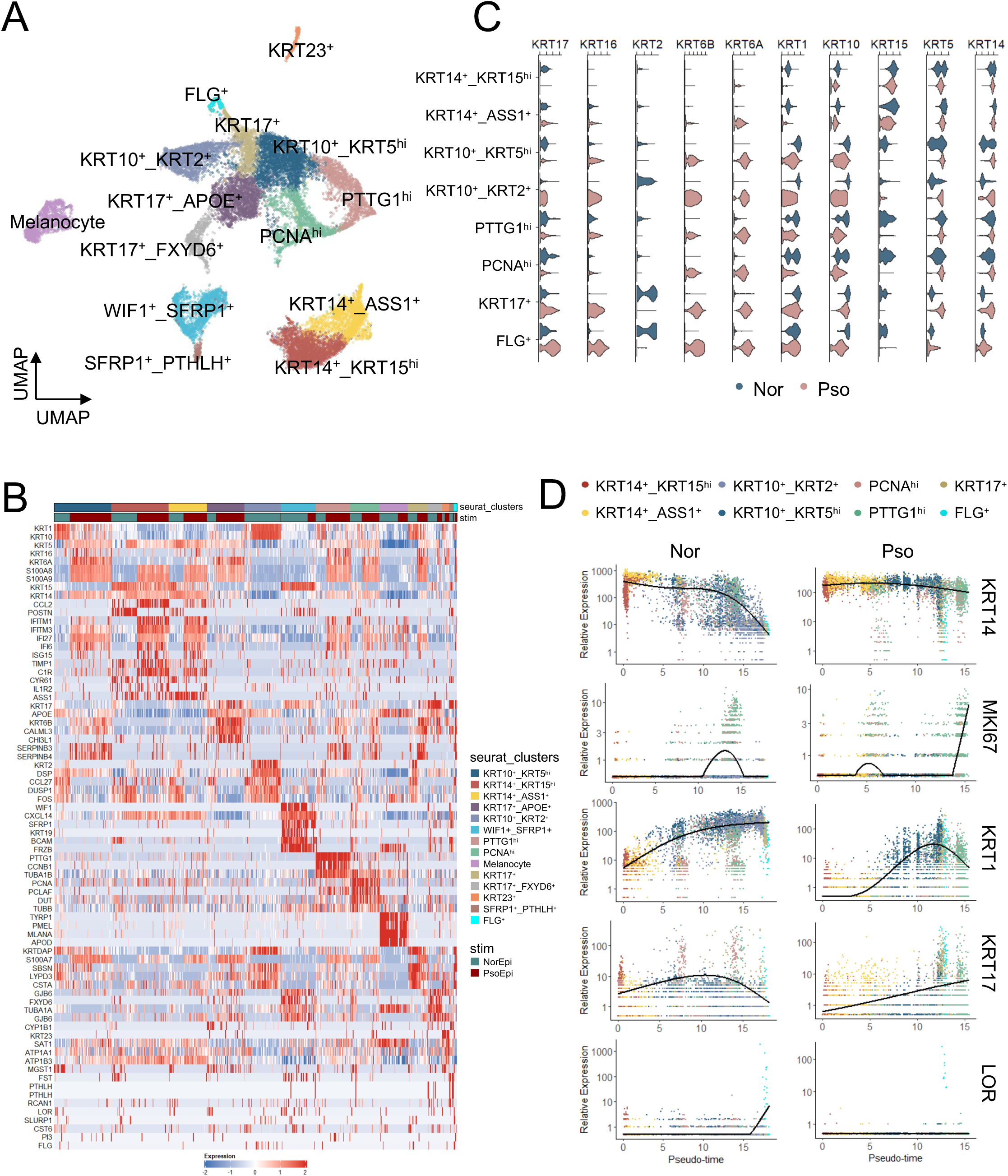
Sub-clustering of keratinocytes in psoriatic and normal skin. (A) UMAP dimensional reduction and sub-clustering of keratinocytes from psoriatic epidermis samples (n = 6) and healthy controls (n = 4). (B) Heatmap of signature genes in keratinocyte sub-clusters shown in (A). (C) KRT genes in keratinocyte sub-clusters of psoriatic epidermis samples and healthy controls. (D) Pseudo-time trajectories of keratinocytes in psoriatic epidermis samples and healthy controls.

**Figure S3.**
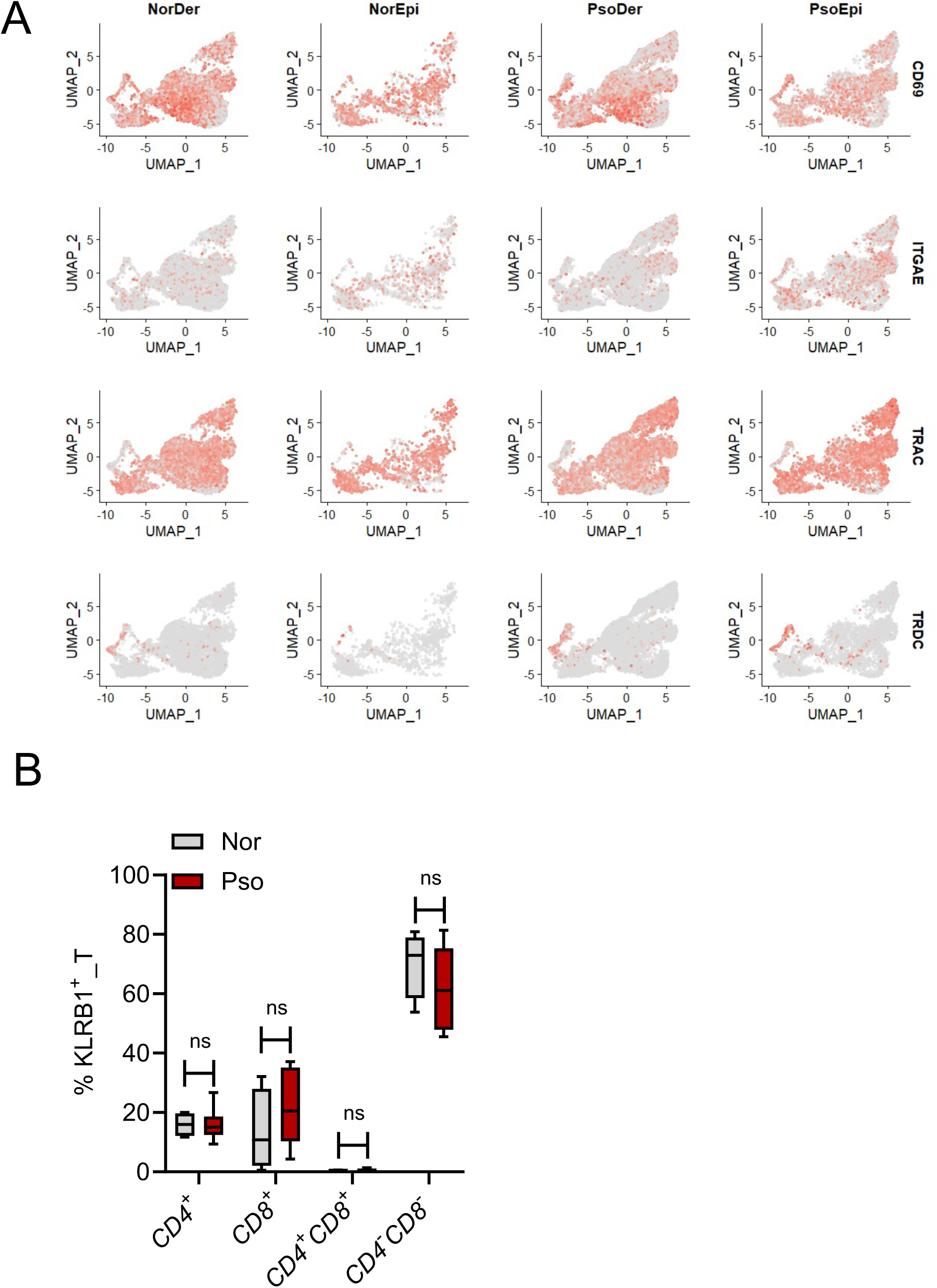
Characterization of *KLRB1*^+^ T cells. (A) Feature-plots of indicated genes in T cells. (B) Percentages of *CD4^+^*, *CD8^+^*, *CD4^+^CD8^+^* and *CD4^-^CD8^-^*T cells in *KLRB1^+^* T cells from psoriatic samples and healthy controls. Data represent the 25^th^ to 75^th^ percentiles (whiskers showing min to max). ns, not significant.

**Figure S4.**
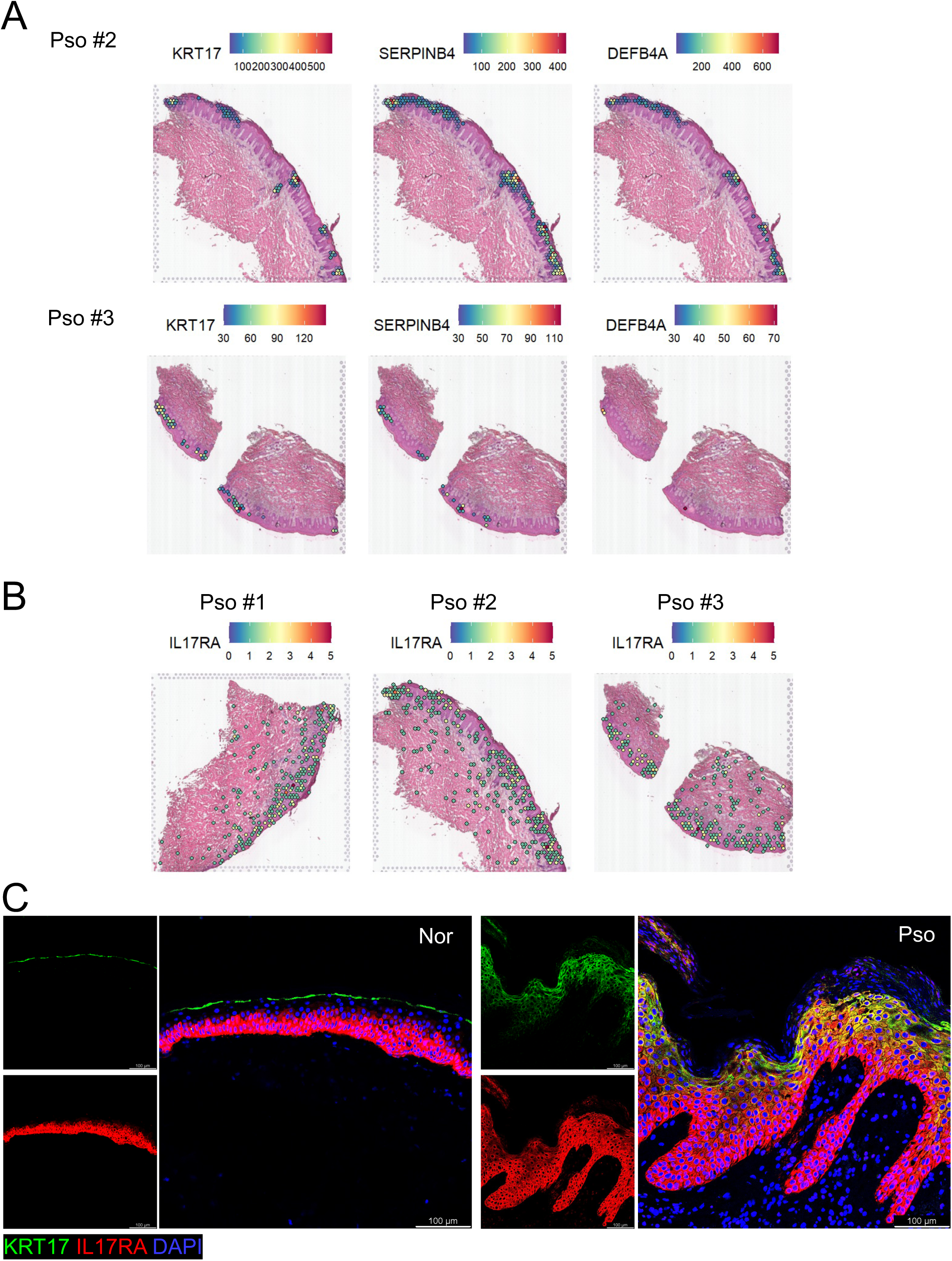
Spatial distribution of IL-17A-downstream signatures. (**A**, **B**) Spatial feature-plots of indicated genes in psoriatic skin sections. (C) Representative immunofluorescence images of KRT17 and IL17RA expression in normal (n = 3) and psoriatic skin sections (n = 5). Data are representative of two independent experiments.

**Figure S5.**
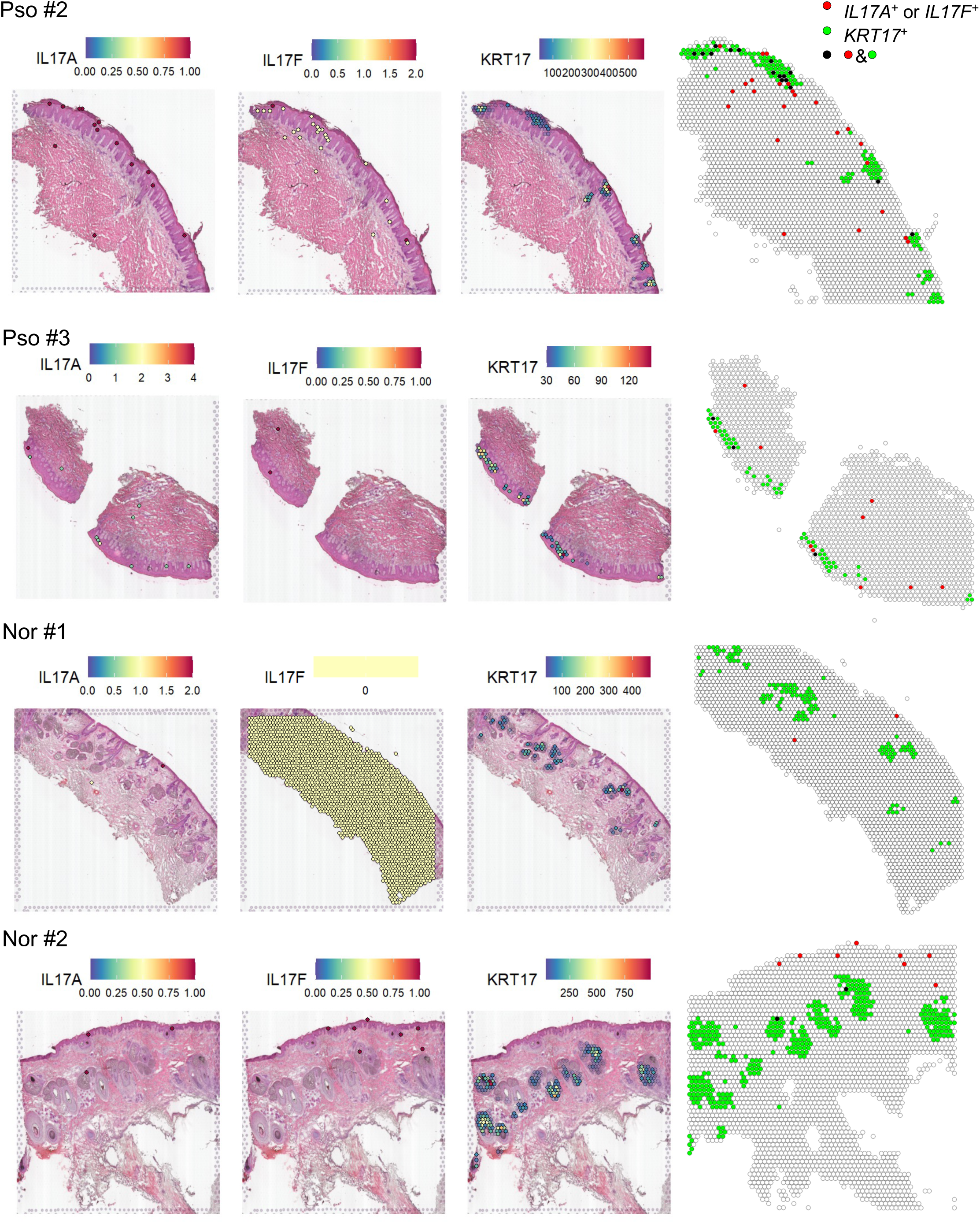
Co-localization of IL-17 and KRT17^+^ keratinocytes in psoriatic epidermis. Spatial feature-plots of *IL17A* and *IL17F* in skin sections from psoriatic patients and healthy donors and merged with KRT17^+^ (*KRT17* expression > 30) spots.

**Figure S6.**
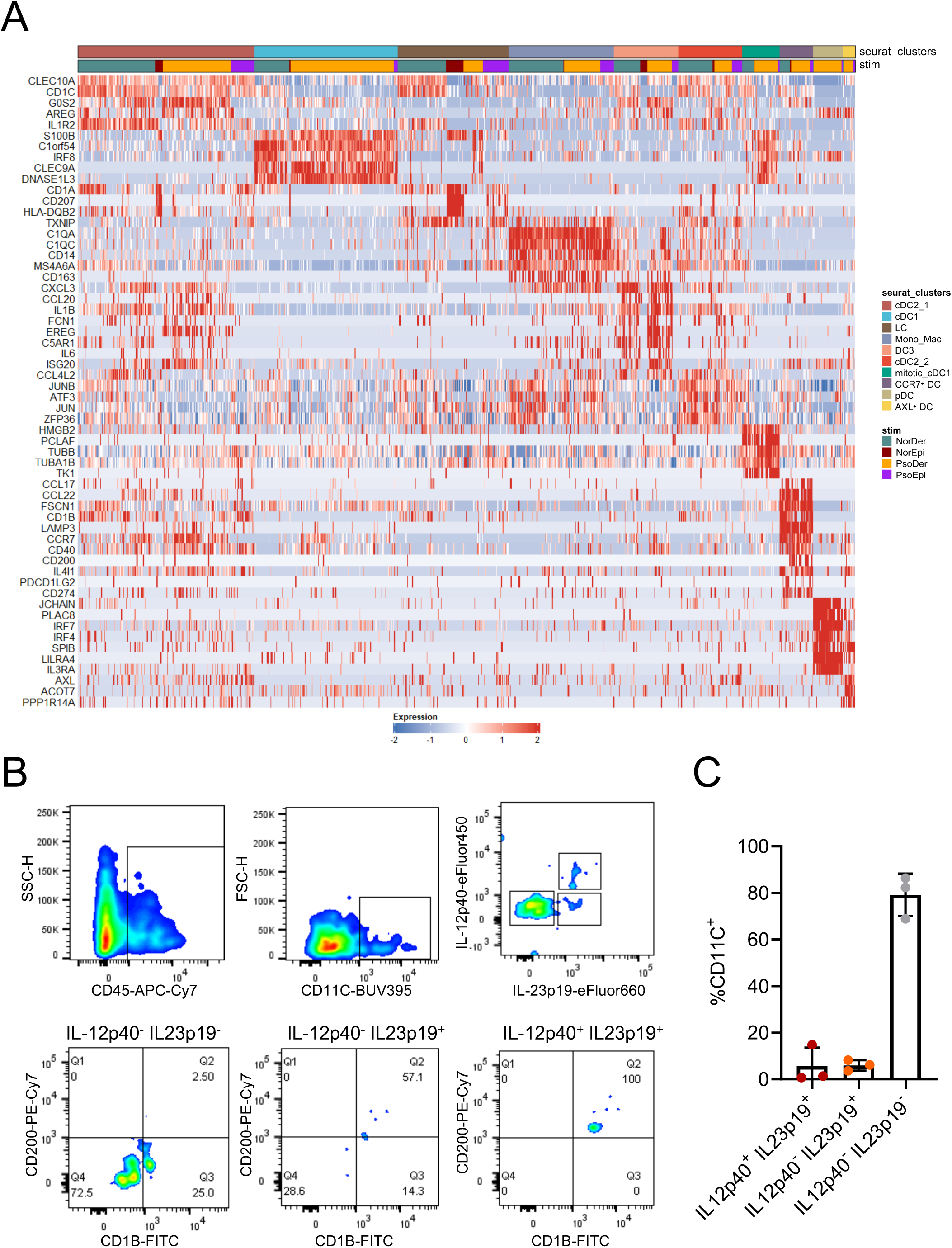
DC and monocyte subtypes in psoriatic and normal skin. (A) Heatmap of signature genes in DC and monocyte subtypes in Figure 3A. (B) Representative flow cytometry plots of IL-12p40^+^IL-23p19^+^, IL-12p40^-^IL-23p19^+^ and IL-12p40^-^IL-23p19^-^ cells in CD45^+^CD11C^+^ cells from psoriatic skin samples (n = 3). (C) Percentages of IL-12p40^+^IL-23p19^+^, IL-12p40^-^IL-23p19^+^ and IL-12p40^-^IL-23p19^-^cells in CD45^+^CD11C^+^ cells from psoriatic skin samples (n = 3). Data represent the mean ± SEM. Data are representative of three independent experiments.

**Figure S7.**
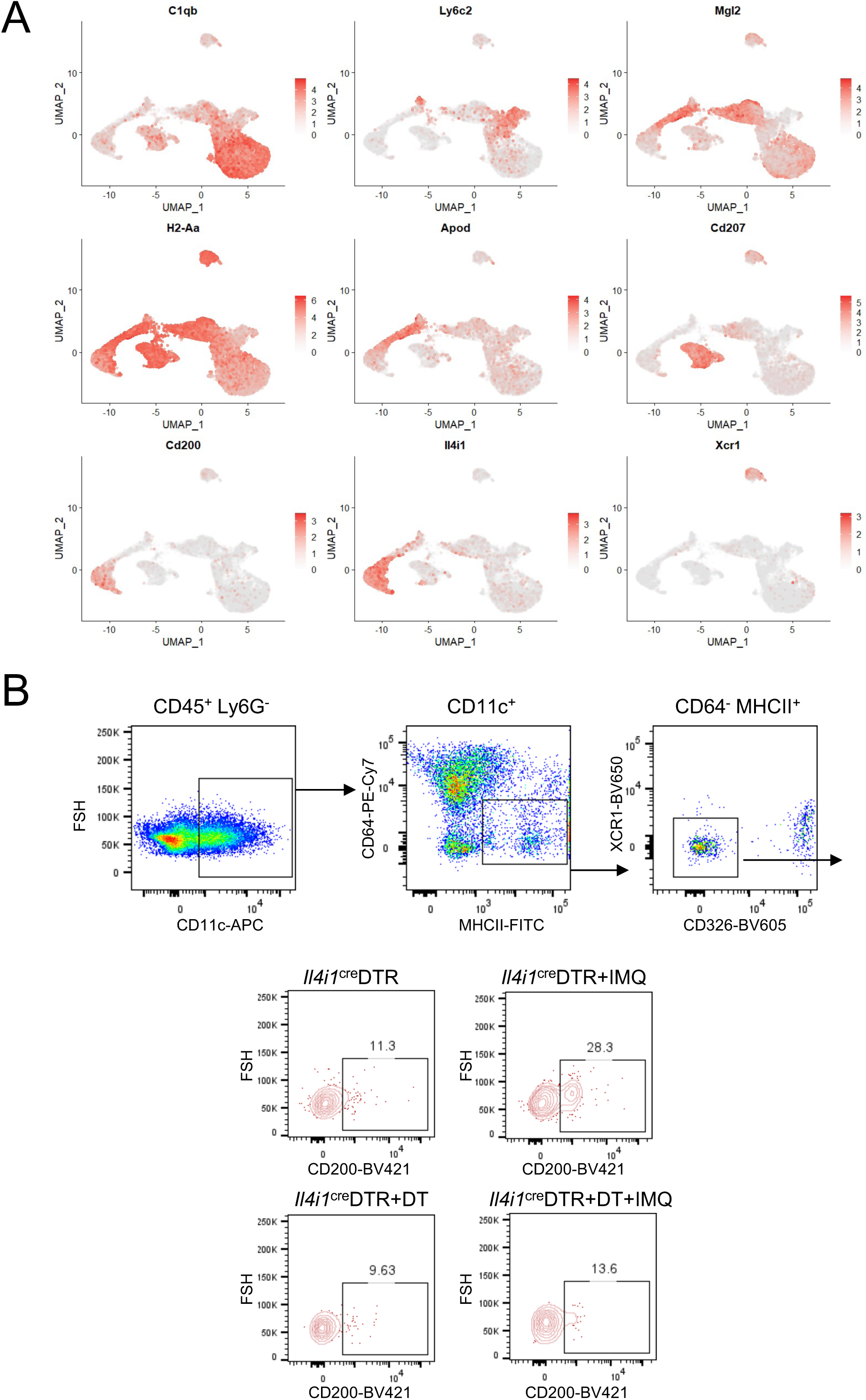
Characterization of myeloid cells in IMQ-induced murine skin. (A) Feature-plots of indicated genes in CD45^+^Ly6G^-^CD3^-^CD19^-^ myeloid cells from mouse ears treated or not with IMQ for 2 days or 5 days. (B) Representative flow cytometry plots of CCR7^+^ DC subsets in cDC2 (CD45^+^Ly6G^-^CD11c^+^MHCII^+^CD64^-^CD326^-^XCR1^-^) of ears from *Il4i1*^cre^DTR mice treated under the indicated conditions (n = 3∼5). Data are representative of two independent experiments.

**Figure S8.**
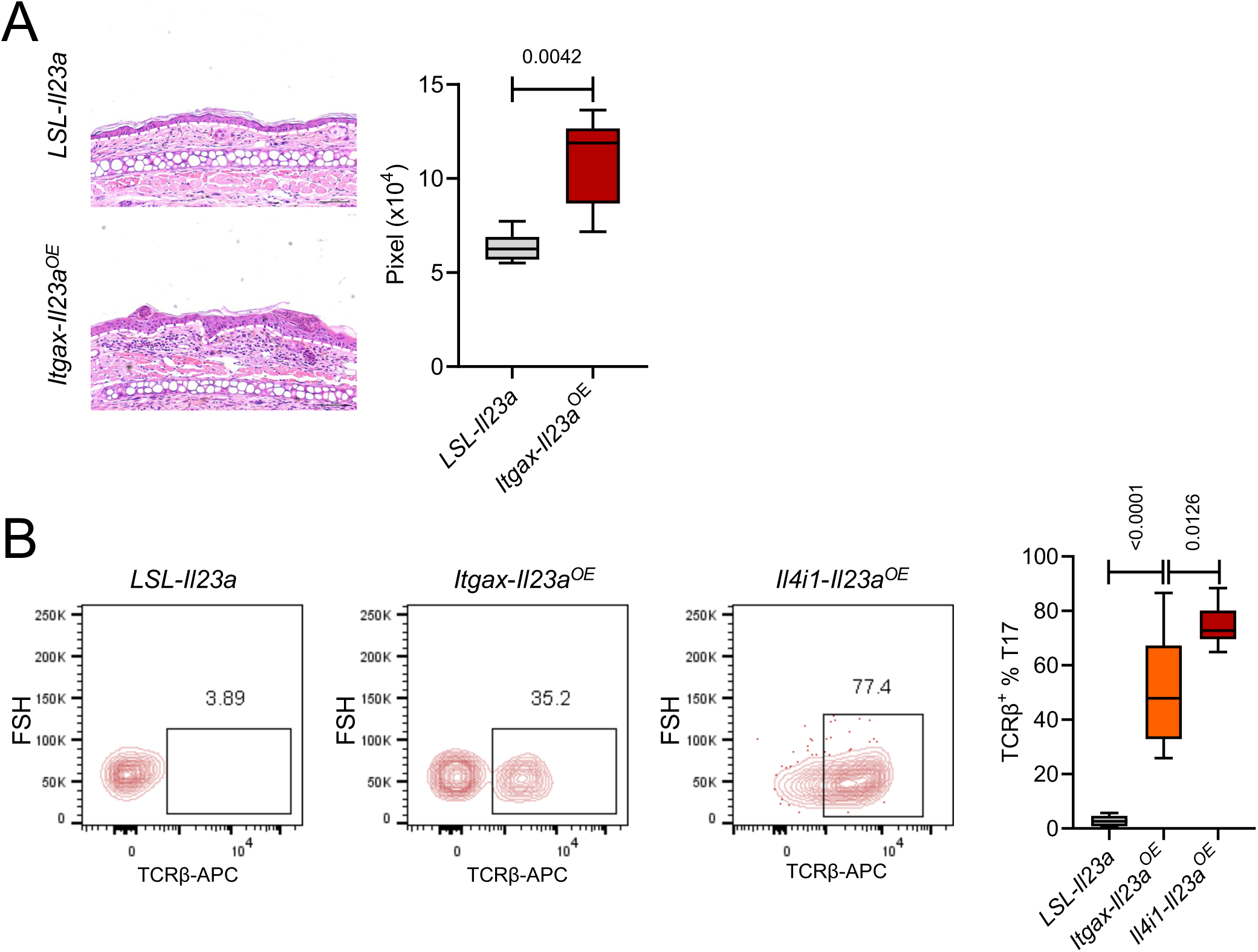
Pathological features of ear skin in *Itgax-Il23a^OE^* mice. (A) Representative H&E images and quantification of acanthosis of 12-week-old *Itgax-Il23a^OE^* mice (n = 5) and *LSL-Il23a* mice (n = 5). Scale bar, 50 μm. Data represent the 25^th^ to 75^th^ percentiles (whiskers showing min to max). *P*-value was determined by two-tailed unpaired *t*-test. (B) Flow cytometry showing the percentages of TCRβ^+^ cells in CD3^+^IL-17a^+^ T cells (T17) of ears from 12-week-old *LSL-Il23a* mice (n = 8), *Itgax-Il23a*^OE^ mice (n = 6) and *Il4i1-Il23a^OE^* mice (n = 6). Data represent the 25^th^ to 75^th^ percentiles (whiskers showing min to max). *P*-values were determined by one-way ANOVA. Data are representative of three independent experiments.

**Figure S9.**
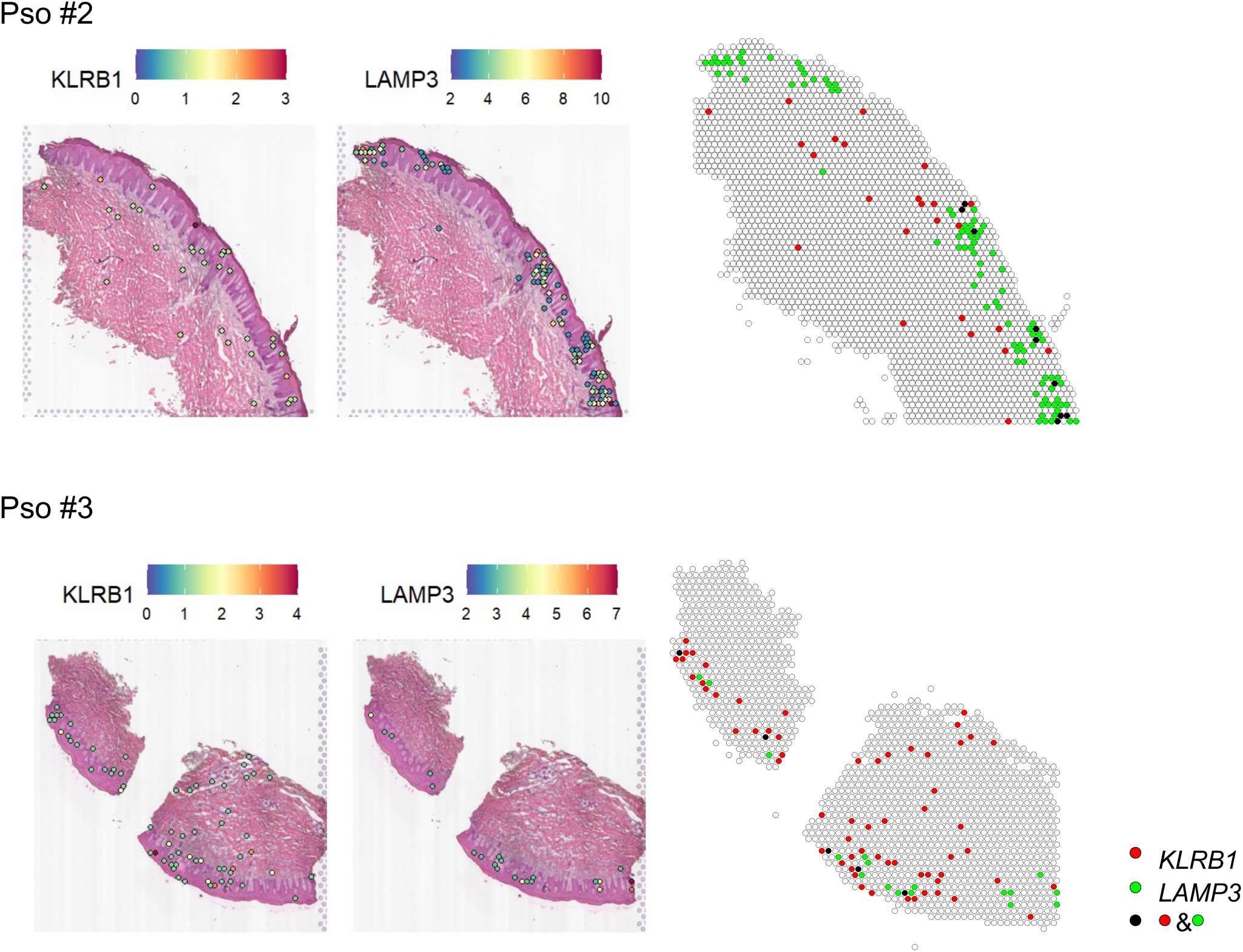
Co-localization of KLRB1 and LAMP3 in psoriatic epidermis. Spatial feature-plots of *KLRB1*and *LAMP3* in skin sections from psoriatic patients and the merged plots.

